# Interaction with AK2A links AIFM1 to cellular energy metabolism

**DOI:** 10.1101/2024.09.09.611957

**Authors:** Robin Alexander Rothemann, Egor Pavlenko, Mrityunjoy Mondal, Sarah Gerlich, Pavel Grobushkin, Sebastian Mostert, Julia Racho, Konstantin Weiss, Dylan Stobbe, Katharina Stillger, Kim Lapacz, Silja Lucia Salscheider, Carmelina Petrungaro, Dan Ehninger, Thi Hoang Duong Nguyen, Jörn Dengjel, Ines Neundorf, Daniele Bano, Simon Pöpsel, Jan Riemer

## Abstract

Apoptosis inducing factor 1 (AIFM1) is a flavoprotein essential for mitochondrial function and biogenesis. Its interaction with MIA40, the central component of the mitochondrial disulfide relay, accounts for some, but not all effects of AIFM1 loss. Our high-confidence AIFM1 interactome revealed novel interaction partners of AIFM1. For one of these interactors, adenylate kinase 2 (AK2), an essential enzyme maintaining cellular adenine nucleotide pools, AIFM1 binding specifically stabilized the isoform AK2A via interaction with its C-terminus. High resolution cryo-EM and biochemical analyses showed that both, MIA40 and AK2A bind AIFM1’s C-terminal β-strand, enhancing NADH oxidoreductase activity by locking an active, dimer conformation and, in the case of MIA40, affecting the cofactor binding site. The AIFM1-AK2A interaction is crucial during respiratory conditions. We further identified ADP/ATP translocases and the ATP synthase as AIFM1 interactors, emphasizing its important regulatory role as a central, organizing platform in energy metabolism.

## INTRODUCTION

Apoptosis inducing factor mitochondrial 1 (AIFM1) is a flavin adenine dinucleotide (FAD)-dependent oxidoreductase of the mitochondrial intermembrane space (IMS). It plays pleiotropic roles in the biogenesis, assembly and maintenance of mitochondrial respiratory chain complexes, in maintaining mitochondrial function and redox control, and in promoting non-caspase-dependent cell death ^1–5^. AIFM1 defects in human patients result in mitochondrial dysfunction associated with neurological symptoms, neurodegenerative diseases, muscle weakness and atrophy, and cardiomyopathy ^4^. In line with these observations, a mouse model with strongly reduced levels of AIFM1, the harlequin (Hq) mouse, suffered for example from progressive cerebellar ataxia, optic tract dysfunction, and hypertrophic cardiomyopathy ^6–8^. Likewise, a knock-in mouse model expressing the first disease-causing *AIFM1* mutation identified in humans ^9^ exhibited early-onset myopathy ^10, 11^. On the molecular level, low protein amounts of AIFM1 in the Hq mouse, in AIFM1 knock-out or depletion cells, strongly reduce complex I subunit levels, and to a lesser extent, complex III and IV subunits of the respiratory chain ^10–15^.

The impact of AIFM1 on respiratory chain complexes can be in part explained by the link between AIFM1 and the mitochondrial disulfide relay system. The import receptor and oxidoreductase MIA40 (in humans also CHCHD4) is the major component of the disulfide relay, responsible for the import and oxidative folding of most soluble IMS proteins, among them structural subunits of complex I and assembly factors of complex I, III, IV, and V ^3, 4, 16–21^. AIFM1 plays a two-fold role in the disulfide relay. Firstly, it facilitates the import of MIA40 itself, and secondly, it forms a stable complex with MIA40 ^13–15, 22, 23^. Notably, the overexpression of MIA40 can complement the loss of AIFM1 in different cell lines, however, not all substrates of the disulfide relay are equally affected by AIFM1 loss ^13–15^.

AIFM1 exhibits a complex NADH-dependent redox behavior and is thought to be capable of sensing NADH levels in the IMS. To fulfil its redox function, AIFM1 possesses two Rossman fold FAD- and NADH-binding domains (**Figure 1A**) ^24^. Binding of NADH leads to the formation of an air-stable FADH_2_-NAD^+^ charge-transfer-complex (CTC) ^25–27^. CTC establishment induces AIFM1 dimerization and the release of a regulatory segment of the C-terminal domain termed C-loop ^25, 28–30^.

**Figure 1.**
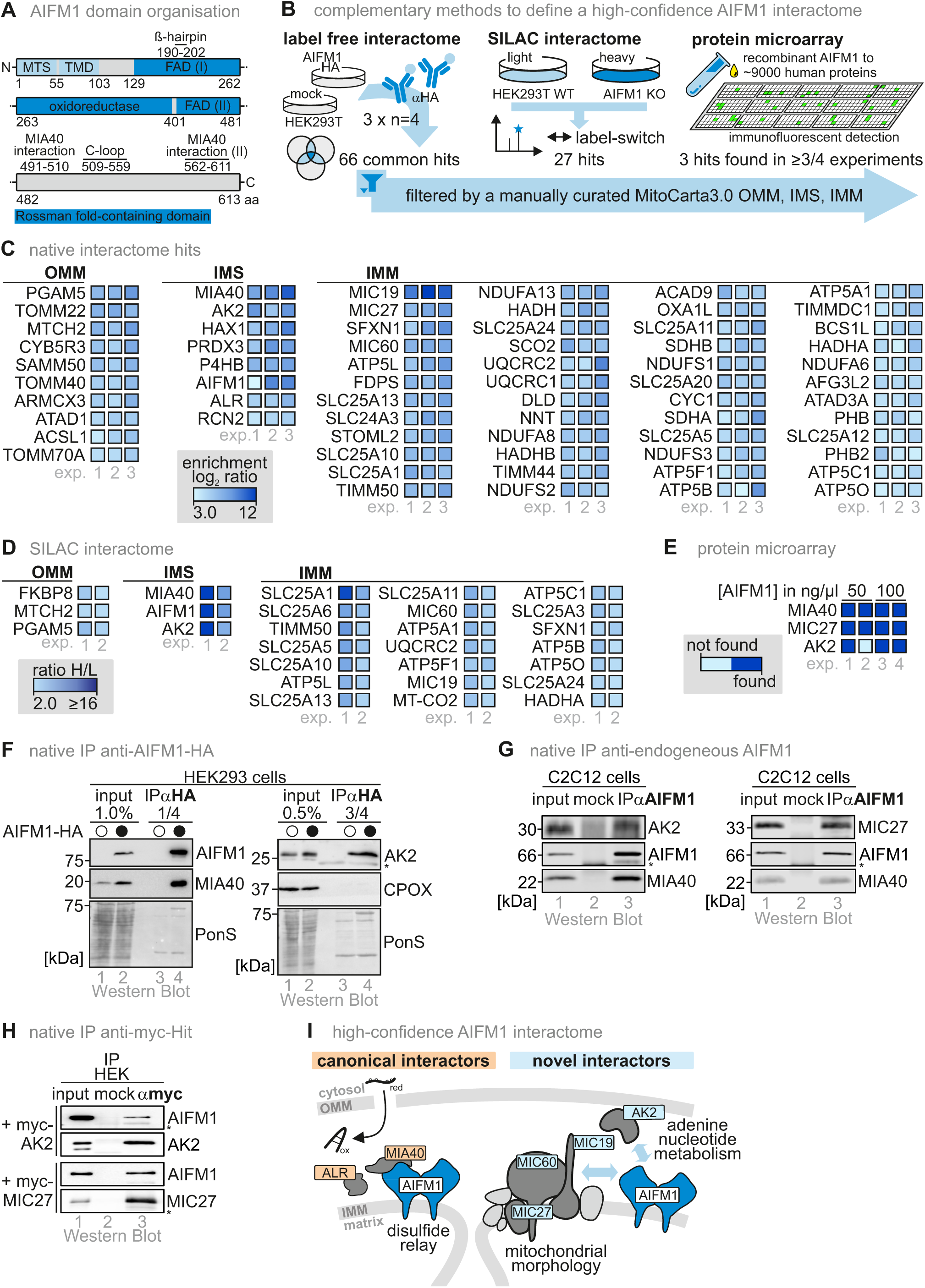
A high-confidence interactome of AIFM1 reveals AK2 and MICOS components as interaction partners. **(A)** Domain layout of AIFM1. The mitochondrial targeting signal (MTS) includes amino acids (aa) 1-55 followed by a sorting transmembrane domain (TMD) until aa 103. The two Rossman folds for FAD- and NADH-binding encompass aa 129–170, 203–261, and 403–479, and 171–202 and 265–399, respectively. The C-loop, includes aa 509-559 and the β-hairpin aa 190-202. **(B)** Approaches to determine a high-confidence interactome of AIFM1. Three different approaches to identify novel interaction partners of AIFM1 were employed: Native immunoprecipitation of AIFM1-HA followed by label-free proteomics, native immunoprecipitation of AIFM1-HA in a SILAC-labeling-based approach, and a native protein-protein profiling microarray approach. **(C)** Interaction partners of AIFM1-HA from native label-free co-immunoprecipitation experiments (IP). Cells were treated with NEM to avoid aberrant disulfide formation, and lysed under native conditions. AIFM1-HA was enriched using HA-antibody beads and eluates were analyzed by label-free mass spectrometry. The experiment was reproduced three times. Each experiment relies on the analysis of 4 biological replicates rendering the results as indicative for 12 co-immunoprecipitation experiments. Fold enrichment in the AIFM1-HA IP was plotted for significant hits of the 3 experimental repeats. A hit is significantly enriched in all three experimental repeats and localizes to IMM, OMM or IMS. N = 12 biological replicates, an unpaired one-sample two-sided Student’s t-test was applied (p<0.2, log_2_-enrichment >3). **(D)** Interaction partners of AIFM1-HA from native SILAC-based co-immunoprecipitation experiments (IP). AIFM1-HA and Mock cells were grown in SILAC medium. Cells were treated with NEM to avoid aberrant disulfide formation, mixed and lysed under native conditions. AIFM1-HA was enriched using HA-antibody beads and eluates were analyzed by label-free mass spectrometry. The experiment was reproduced with inverted isotope labeling. Proteins were counted as hits if at least two peptides per protein were enriched more than 2-fold and the proteins have a localization in the IMS, the IMM or OMM. N = 2 biological replicates. **(E)** Identification of AIFM1 interaction partners by protein-protein microarray. Microarrays coated with ca 9,000 different human proteins were incubated with 50 or 100 ng/µl of recombinant human AIFM1. After washing, bound AIFM1 was detected by immunodecoration. 12 putative AIFM1 interacting partners localizing to IMM, OMM and IMS were detected; 3 of them were common candidates in at least three out of four replicates. N = 4 biological replicates. **(F)** IP of AIFM1-HA and test for interaction with identified potential interaction partners. HEK293 cells expressing AIFM1-HA were treated with NEM and lysed under native conditions, and AIFM1-HA was precipitated using HA-antibody beads. Immunoblot analyses were performed against AK2, MIA40, HA, and as non-interacting control, the IMS protein CPOX. PonS, ponceau staining; asterisk, antibody chains. **(G)** IP of endogenous AIFM1 and test for interaction with identified potential interaction partners. C2C12 cells were lysed under native conditions, and AIFM1 was precipitated using AIFM1-antibody beads. Immunoblot analyses were performed against AK2, MIC27, MIA40, and AIFM1. Asterisk, antibody chains. **(H)** IP of myc-tagged AK2 and MIC27 to test for interaction AIFM1. HEK293 cells were transiently transfected with plasmids expressing myc-AK2 or myc-MIC27. Cells were lysed under native conditions, and myc-AK2 and myc-MIC27 were precipitated using myc-antibody beads. Immunoblot analyses were performed against AK2, MIC27, and AIFM1. Asterisk, antibody chains. **(I)** MICOS subunits and AK2 are confirmed high-confident interaction partners of AIFM1 linking AIFM1 to cellular metabolism and mitochondrial morphology.

The NAD-binding H454 and F310 residues allosterically transmit the structural changes trigged by NADH-binding. Upon their re-arrangement, the AIFM1 dimerization interface is changed to promote dimer formation, and a concerted remodeling of aromatic side chains, also termed the ‘aromatic tunnel’, is transmitted to the C-terminal domain. Ultimately, the resulting movement of the “β-hairpin” releases the C-loop from its binding site and makes accessible a binding pocket that is thought to accommodate electron acceptors of AIFM1 and was shown to be a binding site of a second NADH molecule ^28, 30^. It has been proposed that NADH levels in the IMS under non-stress conditions are sufficient to drive AIFM1 dimerization, while reduced NADH availability for example during starvation may lead to a shift towards monomeric AIFM1 ^1, 15, 29^.

AIFM1 dimerization is critical for its function. For example, only the dimer facilitates MIA40 import and binding ^13, 15, 23, 31^. Biochemical and structural approaches defined the MIA40-AIFM1 interaction interface. In MIA40, the mostly unstructured N-terminal region is critical for interaction ^13, 15^. The recent crystal structure of AIFM1 in complex with an N-terminal MIA40 peptide revealed the interaction of a MIA40 β-hairpin (amino acids, aa 3-15) with the AIFM1 C-terminal region (aa 491-510, 562-611) by parallel β-strand complementation ^31^. These studies have suggested that, in order to establish the MIA40 binding site, C-loop displacement is required.

Disease-causing AIFM1 variants lead to AIFM1 deficiency and heterogeneous clinical manifestations in which the main affected tissues exhibit aberrant mitochondrial oxidative phosphorylation (OXPHOS) ^1, 4, 20, 32^. Given the evident pleiotropic effects of pathogenic AIFM1 mutations, we reasoned that AIFM1 deficiency may lead to variable metabolic phenotypes by altering the localization and/or activity of other binding partners. To test this hypothesis, we employed complementary biochemical, structural and cell biology analyses to acquire a high-confidence interactome of AIFM1, and to understand structural and functional implications of these novel interactions with AIFM1. We identified adenylate kinase 2 isoform A (AK2A) and components of the mitochondrial contact site and cristae organizing system (MICOS) as AIFM1 interaction partners. The MICOS complex is important to establish and maintain the delicate mitochondrial morphology. AK2 plays an important role in maintaining balanced adenine nucleotide pools in the IMS by catalyzing the reversible transfer of phosphate groups among adenine nucleotides (2ADP ↔ AMP + ATP). This activity aids in shuttling ATP/ADP/AMP across the IMS, and in establishing ATP/ADP gradients across the IMM to allow exchange of these nucleotides by the mitochondrial ADP/ATP carriers (SLC25A4, SLC25A5, SLC25A6) ^33–36^. Notably, in organs with high energy demand such as the heart, AK2 appears to account for almost half of the total cellular adenylate kinase activity emphasizing its critical role in respiration ^37, 38^. The loss of AK2 leads to impaired mitochondrial function ^39, 40^, hampers induction of the endoplasmic reticulum unfolded protein response ^39^, and sensitizes cells to induction of apoptosis ^41–43^. AK2 knockout in mice is embryonic lethal (E7.5) ^44^, while human patients with defects in AK2 suffer from an autosomal recessive form of severe combined immunodeficiency named reticular dysgenesis ^45, 46^.

To structurally characterize the AIFM1-AK2A interaction that we found to be stable and dependent on AIFM1 dimerisation, we obtained high resolution structures of the AIFM1 dimer alone or bound to MIA40 or AK2A using single-particle cryo-electron microscopy (cryo-EM). MIA40 and AK2 compete for the same interaction interface in AIFM1, and both interaction partners bind via parallel β-strand complementation. Binding of either MIA40 or AK2A has two consequences: stabilization of the AIFM1 dimer and increased redox activity towards an artificial electron acceptor, enabling NADH-oxidizing activity of AIFM1 at close to the physiological NADH concentration. Lastly, we reveal a potential function of the AIFM1-AK2A interaction in supporting the metabolic switch from fermentative carbon sources to carbon sources that require respiration. Our results support the notion that this is due to a positioning of AK2 close to ADP/ATP translocases in the IMM, which we demonstrate by different AIFM1 interaction studies.

## RESULTS

### A high-confidence interactome reveals AK2 and MICOS components as novel AIFM1 interaction partners

To investigate which proteins are influenced by AIFM1, we explored the interactome of AIFM1. Since AIFM1 is a membrane protein bound to the IMM in the crowded environment of the IMS, we employed complementary approaches for the identification of AIFM1 interaction partners to obtain a high confidence interactome (**Figure 1B**).

First, we performed native immunoprecipitation from HEK293 cells stably expressing C-terminally HA-tagged AIFM1 or from mock cells, followed by quantitative label-free proteomic analysis. We performed three independent sets of experiments with four biological replicates each, and identified 66 proteins with IMS localization or domains facing the IMS as potential interaction partners of AIFM1 (**Figures 1B** and **C Supplementary Figure S1A**). We then employed a similar but SILAC (stable isotope labeling with amino acids in cell culture)-based approach, which allows a precise quantification of enrichment (**Figures 1B** and **D, Supplementary Figure S1B**). We identified 27 proteins with IMS localization or domains facing the IMS as potential interaction partners of AIFM1. We complemented these analyses by an unbiased *in vitro* protein-protein profiling of putative AIFM1 interactors. We employed microchips containing purified recombinant human proteins and incubated them with purified AIFM1 (soluble AIFM1 lacking the transmembrane domain, ^24^). Subsequent decoration of the microarrays with an AIFM1 antibody revealed three proteins that fulfilled our stringent selection criteria for a potential interaction with AIFM1 *in vitro* (**Figures 1B** and **E, Supplementary Figure S1C**).

By integrating these data sets, we retrieved the proteins MIA40, AK2, and MICOS components (MIC27 in all three approaches; MIC19 and MIC60 in two approaches) as common AIFM1 interactors. MIA40 has previously been identified to stably interact with AIFM1 ^13–15, 22^, supporting the validity of our approaches. We verified interaction partners by immunoprecipitation of AIFM1-HA from stable inducible HEK293 cells (**Figure 1F**). Immunoprecipitation of endogenous AIFM1 from C2C12 cells and subsequent immunoblotting against MIA40, AK2 and MIC27 further confirmed the interaction (**Figure 1G**). Lastly, immunoprecipitation of myc-AK2 and myc-MIC27 expressed in HEK293 cells and subsequent immunoblotting against AIFM1 confirmed their interaction in a reciprocal manner (**Figure 1H**). MIA40 served as positive control in all approaches. Together, we identified AK2 and MICOS components (MIC27, MIC19, MIC60) as high-confidence interaction partners of AIFM1, thus establishing its link to energy metabolism, and the formation and maintenance of mitochondrial morphology (**Figure 1I**).

### The C-terminal region in the isoform AK2A is important for AIFM1 interaction

To directly investigate AIFM1 complexes, we performed gel filtration assays and found that AIFM1 formed complexes with an apparent molecular weight (MW) between 100 and 200 kDa. In HEK293 cells, almost the entire share of cellular AIFM1 as well as MIA40 were found in this complex (**Figure 2A**, ^15^). Deletion of AIFM1 resulted in a shift of MIA40 to lower MW fractions (**Figure 2A**, ^15^). Since we expected a similar MW shift for other AIFM1 interaction partners, we performed the gel filtration analysis with wildtype (WT) and AIFM1 knock-out (AIFM1 KO) cells, collected the fractions corresponding to the high MW AIFM1 complex, and subjected them to proteomic analysis (**Figure 2A**). AIFM1, MIA40, UQCRB, COX7B, and AK2 were detected in the AIFM1 complex fractions in the WT but not the AIFM1 KO cell lysates (**Figures 2B**). The identification of AIFM1 and MIA40 as positive controls confirmed the validity of the approach. Importantly, this complementary approach again points to a specific interaction between AIFM1 and AK2 in a stable complex that persists during gel filtration. We further confirmed the AIFM1-AK2 complex in WT, AIFM1 KO, and AIFM1 KO cells complemented with AIFM1-HA using gel filtration and immunoblotting (**Figure 2C**) and in different cell lines (**Supplementary Figure S2A**). This complex also resists cycloheximide treatment for 4 hours indicating that once formed the complex is stable and long-lived similar to the AIFM1-MIA40 complex (**Supplementary Figure S2B**).

**Figure 2.**
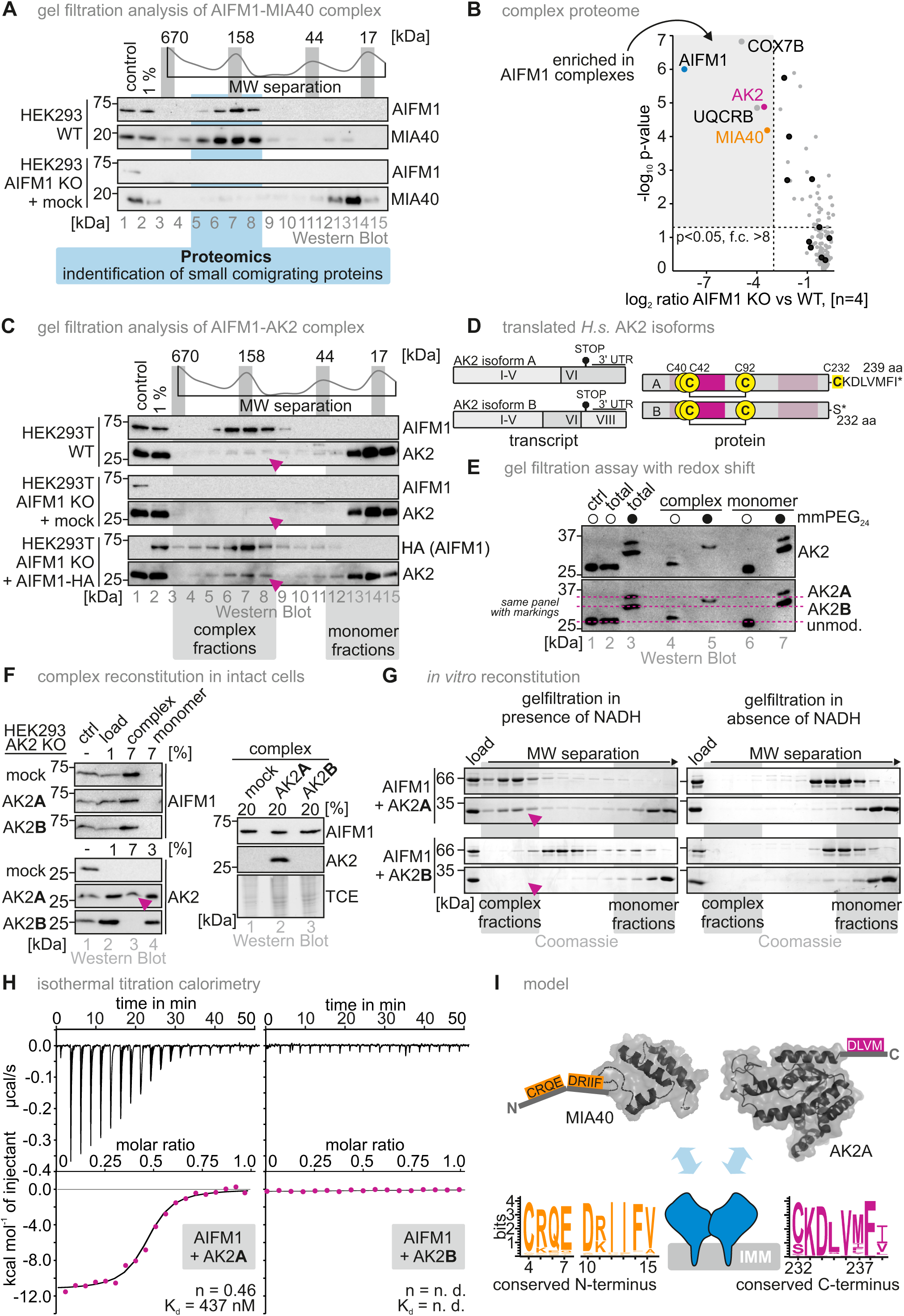
The isoform AK2A forms via its conserved C-terminus a stable complex with the AIFM1 dimer. **(A)** Proteomic analysis assessing proteins depleted of the AIFM1-MIA40 complex fractions in AIFM1 KO cells. WT HEK293 cells, and cells lacking AIFM1 (AIFM1 knockout, AIFM1 KO) were lysed under native conditions (Triton X-100), and the cleared lysates were subjected to gel filtration analysis. Eluted fractions were subjected to TCA precipitation, resuspended in loading buffer containing SDS and DTT, and analysed by immunoblotting. Endogenous AIFM1 and endogenous MIA40 migrated in a complex with a size of around 150 kDa (as judged by comparison to protein markers: thyroglobulin, 670 kDa; γ-globulin, 158 kDa; ovalbumin, 44 kDa; myoglobulin, 17 kDa). Absence of AIFM1 resulted in the migration of MIA40 at the height of monomeric MIA40. Fractions of the complex region (light blue) were submitted to quantitative label-free proteomic analysis. **(B)** Results of the gel filtration-coupled proteomics experiment. As expected, MIA40 and AIFM1 are strongly depleted in AIFM1 KO cells at the height of the complex. Applying stringent parameters (fold change >8) results in the identification of three additional proteins that might potentially reside in the AIFM1-MIA40 complex, AK2, COX7B and UQCRB. N = 4 biological replicates, an unpaired one-sample two-sided Student’s t-test was applied (p<0.05, fold change >8). **(C)** Gel filtration analysis of WT, AIFM1 KO and AIFM1 KO cells complemented with AIFM1-HA HEK293 cells. Experiment was performed as described in **(A)**. The majority of the highly abundant AK2 migrated at its monomeric mass. About 5% of the protein was found reproducibly in the same fractions as the MIA40-AIFM1 complex (purple arrowhead). This higher molecular weight fraction of AK2 disappeared upon loss of AIFM1 and reappeared after reintroduction of AIFM1-HA. For better representation, we combined in subsequent experiments often complex and monomer fractions. **(D)** Human AK2 is present in multiple splice forms. Most prominent are AK2A and AK2B that as proteins only differ in the last 8 amino acids. The remainder of the proteins contain the important sites for activity (P loop, lid and NMP-binding domain), and also the structural disulphide bond. AK2B stops after 232 amino acid residues with S232, while AK2A stops after 239 amino acids. Instead of S232, AK2A bears C232. This additional cysteine residue in AK2A (total of 4 cysteines) compared to AK2B (3 cysteines) can aid in distinguishing these two very similar isoforms of the protein. **(E)** Gel filtration assay of HEK293 cells coupled to mmPEG shift assay. Experiment was performed as described in **(A)** but instead of the proteomic analysis, fractions were incubated with the reductant TCEP and the thiol-modifying agent mmPEG24, and subsequently analysed by SDS-PAGE and immunoblotting. The treatment with mmPEG24 modifies all four cysteines in AK2A and all three cysteines in AK2B. The modification of one additional cysteine in AK2A leads to slower migration of AK2A on SDS-PAGE. The majority of cellular AK2 appears to be AK2B but it is AK2A that comigrates with AIFM1 in the higher molecular weight complex. Complex and monomer fractions are presented. **(F)** Gel filtration assay of HEK293 cells depleted of AK2 (AK2 knockout, AK2 KO) complemented with an empty vector (Mock), AK2A or AK2B. Experiment was performed as described in **(A)** but instead of the proteomic analysis, fractions of the region containing monomeric AK2 and the region containing AK2 in complex were separately pooled, TCA precipitated and analysed by SDS-PAGE and immunoblotting. Only AK2A can be detected in the complex fraction. **(G)** *In vitro* reconstitution of the AK2A-AIFM1 complex and analysis by gel filtration. Purified AIFM1 was incubated with purified AK2A or AK2B in the presence or absence of NADH. Formation of a higher molecular weight (MW) complex of AIFM1 and AK2 is only observable in the presence of NADH when AK2A is used. Since NADH is required for dimerization of AIFM1 this implies that like for the MIA40-AIFM1 complex only the AIFM1 dimer can bind to AK2A. **(H)** Isothermal Titration Calorimetry (ITC) analysis of AIFM1 together with AK2A or AK2B in the presence of NADH. AIFM1 and AK2A bind to each other with a K_d_ of approximately 0.44 μM and stoichiometry between AIFM1 and AK2A of ca. 2 to 1. AK2B does not interact with AIFM1 in this experiment. **(I)** The interaction sites in MIA40 and AK2A for their interaction with AIFM1. The interaction site in MIA40 spans roughly the first 15 amino acids and contains conserved charged and aromatic amino acids. AK2A interacts with AIFM1 via the last 9 amino acids. Likewise, in this amino acid patch conserved charged and aromatic/aliphatic amino acids are present.

Notably, this gel filtration approach is unsuitable for investigating MICOS-AIFM1 interactions as the >700 kDa MW of a mature MICOS-AIFM1 complex would prevent the detection of the relatively small shift upon loss of AIFM1 from this complex. Given the impact of AIFM1 deletion on cellular metabolism and the robust identification of AK2 as AIFM1 interaction partner we performed a detailed mechanistic analysis of the AIFM1-AK2 interaction.

In human cells, the important AK2 isoforms, AK2A and AK2B, differ only in their C-terminal residues. While AK2A is 239 aa in length, the shorter AK2B ends at S232 (**Figure 2D**). These differences do not extend to the enzymatically active domains, and consequently, the purified proteins have similar activities (**Supplementary Figure S3**). Due to their very similar size, it is difficult to experimentally distinguish both AK2 isoforms. Therefore, we made use of a cysteine in AK2A (C232) that is absent in AK2B to differentiate between the isoforms and performed a maleimide shift assay of the endogenous AK2 comigrating with AIFM1 during gel filtration. In this assay, the covalent attachment of the maleimide mmPEG24 to free cysteines is detectable as a shift in apparent MW in SDS-PAGE. Thus, AK2A (four cysteines) can be distinguished from AK2B (three cysteines). Using this assay, we detected only the AK2A isoform in the high MW fractions upon gel filtration (**Figure 2E**). Confirming this finding, in AK2 KO cells expressing AK2A or AK2B, only AK2A but not AK2B interacted with AIFM1 as demonstrated by gel filtration (**Figure 2F**).

We next tested the direct interaction between AIFM1 and AK2 isoforms *in vitro* using recombinant purified AK2A and B and AIFM1 (**Supplementary Figures S3** and **S4**). Gel filtration showed that only AK2A but not AK2B comigrated with AIFM1 and that this interaction was dependent on NADH, suggesting that, similar to the AIFM1-MIA40 interaction ^13, 15, 31, 47^, AK2A binding depends on AIFM1 dimerization induced by NADH (**Figure 2G** and **Supplementary Figure S4**). We then assessed binding between purified AK2 isoforms and AIFM1 using isothermal calorimetry (ITC), confirming that only AK2A interacts with AIFM1 (**Figure 2H**). The ITC measurement showed a K_d_ of 437 nM and a binding stoichiometry of 0.46 indicating that one AK2A molecule interacts with two AIFM1 molecules. The binding is thus slightly weaker compared to MIA40 (K_d_ = 180 nM) but adhered to the same stoichiometry ^15^. Collectively, our findings show that the isoform AK2A interacts with AIFM1 and that the last seven aa, forming a conserved region among AK2A homologs, mediate this interaction (conserved are in particular K233, D234, V236, and F238, **Figure 2I**).

### Structures of the AIFM1-AK2A and AIFM1-MIA40 complexes reveal a shared binding site

To understand the structural basis and implications of AK2A and MIA40 binding, we solved three structures of human soluble AIFM1 (aa 103-613) by cryo-EM: the dimer (1) without an interaction partner (termed ‘AIFM1 dimer’ here), (2) bound to AK2A or (3) bound to MIA40 (**Supplementary Figure S5**). We obtained reconstructions for the AIFM1 dimer, AIFM1-AK2A and AIFM-MIA40 complexes at a global resolution of 2.8 Å, 2.6 Å, and 2.4 Å, respectively (**Supplementary Figures S6-S9**, **Table 1**). We built atomic models of AIFM1 aa 128-611 (lacking aa 511-557 (AIFM1 dimer or AIFM1-MIA40) or aa 511-549 (AIFM1-AK2A) due to flexibility), as well as the AIFM1 interacting residues 232-239 of AK2A and 2-20 of MIA40 into these maps. Despite the use of full-length AK2A and MIA40, only these parts of AK2A and MIA40 were observed in our cryo-EM reconstructions in addition to the AIFM1 dimer, indicating that these are the major, stably interacting residues. Our high-resolution cryo-EM reconstructions of both the non-bound as well as complexed AIFM1 enabled us to analyze the structural implications of MIA40 and AK2A binding in solution with high confidence and detail.

Both AK2A and MIA40 bind to the β-sheet of the AIFM1 C-terminal domain (aa 480-510, 559-580) via parallel β-strand complementation, adding one and two β-strands, respectively (**Figures 3A** and **B**). In the crystal structures of the murine and human AIFM1 dimers, aa 538-544 or aa 510-515, respectively, occupied the same site on one or both of the protomers ^24, 26, 31^. In our reconstructions, clear densities corresponding to F238 of AK2A and F14 of MIA40 show that the added β-strands originate from these proteins rather than AIFM1 (**Supplementary Figures S10A** and **B**). AK2A and MIA40 models were unambiguously built bound to both AIFM1 protomers, indicating that two AK2A or MIA40 molecules can bind to one AIFM1 dimer in solution. We did not obtain reconstructions with only one binding site occupied by either AK2A or MIA40. Possibly, a stabilizing effect of MIA40 and AK2A on the complex resulted in an over-representation of particles corresponding to two MIA40 or AK2A molecules per AIFM1 dimer yielding the best resolved reconstructions.

**Figure 3.**
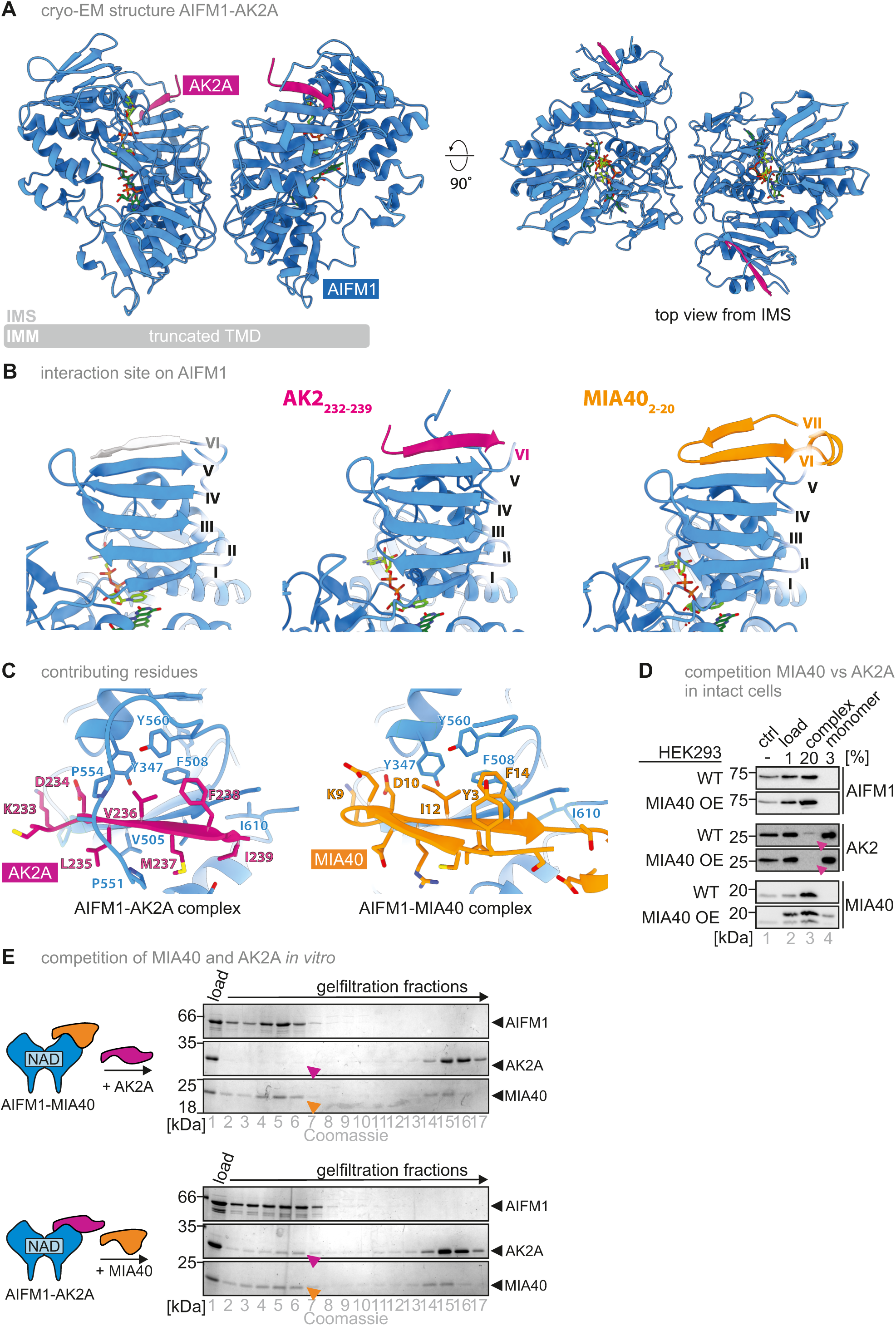
Atomic models of AIFM1 dimer, the AIFM1-AK2A, and the AIFM1-MIA40 complexes reveal binding interfaces. **(A)** Atomic model of the AIFM1 dimer (blue) bound to AK2A (purple). Of AK2A, high-resolution density accounting for aa 232-239 was obtained at both C-terminal domains of the AIFM1 protomers. The remainder of AK2A was presumably flexible and therefore not reconstructed by single-particle cryo-EM at high resolution. **(B)** AK2A (aa 232-239) (purple) and MIA40 (aa 2-20) (orange) bind to the AIFM1 C-terminal domain via parallel β-strand complementation. Grey: additional β-strand of AIFM1 (aa 511-516) in the AIFM1 dimer in the absence of AK2A or MIA40. No densities that could correspond to other parts of AK2A and MIA40 could be detected by cryo-EM nor by negative-stain EM (data not shown), suggesting that the rest of these interaction partners does not stably interact with AIFM1 under the conditions used, but remains largely flexible relative to AIMF1. aa 580-612 of AIFM1 are omitted for clarity. **(C)** Detailed view of residues stabilizing the interaction of AK2A and MIA40 with AIFM1. **(D)** Overexpression of MIA40 in HEK293 cells replaces AK2A from the AIFM1-AK2A complex. Experiment was performed as described in **Figure 2F** with WT HEK293 cells and cells overexpressing MIA40 (MIA40 OE). Purple arrow heads indicate that in MIA40 OE cells, AK2A is lost from the complex fraction. **(E)** *In vitro* competition assay between AK2A and MIA40 for binding to AIFM1. Experiment was performed as described in **Figure 2G** except that after pre-binding of MIA40 to AIFM1, AK2A was added (*upper panel*) or *vice versa* (*lower panel*). While AK2A is not able to bind to AIFM1 if MIA40 is already present, MIA40 can bind to AIFM1 even if AK2A was pre-bound. Purple and orange arrow heads indicate the position of AK2A and MIA40, respectively in the complex fraction.

Two sets of hydrophobic interactions stabilize AK2A and MIA40 binding. On one side of the β-sheet, L235 and M237 of AK2A or I13 of MIA40 interact with V505 and V507 of AIFM1. On the other side, three aromatic residues of AIFM1 (Y347, F508 and Y560) provide a hydrophobic patch that is occupied by V236 and F238 of AK2A or I12 and F14 of MIA40 (**Figure 3C**). The second β-strand of MIA40 is stabilized by Y3 stacking with F14. In addition, conserved Lys and Asp residues, i.e. K233 and D234 of AK2A and K9 and D10 of MIA40, at the C-terminal end of the complementing β-strand form a possible hydrogen bonding network with S500, the backbone carbonyl of L502, and T504 side chain of AIFM1 (**Supplementary Figure S10C**). While MIA40 adds two strands to the AIFM1 β-sheet, AK2A adds only one but additionally binds to a part of the otherwise flexible C-loop (aa 550-558) that traverses the AK2A β-strand, possibly forming hydrogen bonds between the backbone carbonyl of AIFM1 Q525 and the backbone amine of AK2A V236 (**Supplementary Figure S10C**). A similar engagement of AIFM1 aa 550-558 was also reported for a human AIFM1 dimer resolved by X-ray crystallography ^28^. Thus, while the interaction site and critical residues involved are identical for AK2A and MIA40, other aspects of the interaction are divergent. This divergence is also reflected by the differences in binding affinity between MIA40 and AK2A to the AIFM1 dimer (180 nM ^15, 47^ vs. 437 nM (**Figure 2H**), respectively), and aligns well with *in vitro* and *in cell* competition experiments, in which MIA40 was able to replace AK2A partially from the preformed AIFM1-AK2A complex while AK2A could not release MIA40 from the AIFM1-MIA40 complex (**Figure 3D** and **E**). This might imply that under conditions of limited AIFM1 availability MIA40 might outcompete AK2A for binding to AIFM1.

### Structural changes in AIFM1 upon AK2A or MIA40 binding change AIFM1 dimer stability and NADH oxidase activity

To analyze the structural impact of AK2A and MIA40 binding on AIFM1, we first assessed the structural variability between AIFM1 protomers within each of the AIFM1 complexes, aligned to the N-terminal domain near the dimer interface. The AIFM1 dimer and in particular AIFM1-MIA40 show inter-protomer variability predominantly of the NAD binding and parts of the C-terminal domain, whereas little displacement was seen when comparing the AK2A bound protomers (**Supplementary Figure 11**). This indicates that AK2A reduces the inherent structural variability within the AIFM1 dimer. Comparing the conformations of AIFM1 protomers between complexes, AIFM1-AK2A protomers overall closely resembled AIFM1 dimer protomers (**Supplementary Figures S12A** and **B**). MIA40, however, induced up to 3 Å Cα rmsd variability compared to the AIFM1 dimer, particularly of the NAD binding domain, with the strongest displacement seen for the α-helix contacting the C-terminal domain (aa 345-359) (**Supplementary Figure S12C**). The NAD binding domain shifts towards the N-terminal domain and dimer interface, also reflected in the relative position of the NAD and FAD cofactors upon MIA40 binding. Only in AIFM1-MIA40, the distance between the nicotinamide and isoalloxazine rings is reduced by approx. 0.2 Å. The position of the adenines differs by up to > 2 Å comparing protomers from the AIFM1 dimer and the AIFM1-MIA40 complex (**Figure 4A**). Thus, the tight interaction of MIA40 leads to a compaction of AIFM1 that affects the active site via displacement of the NAD binding domain.

**Figure 4.**
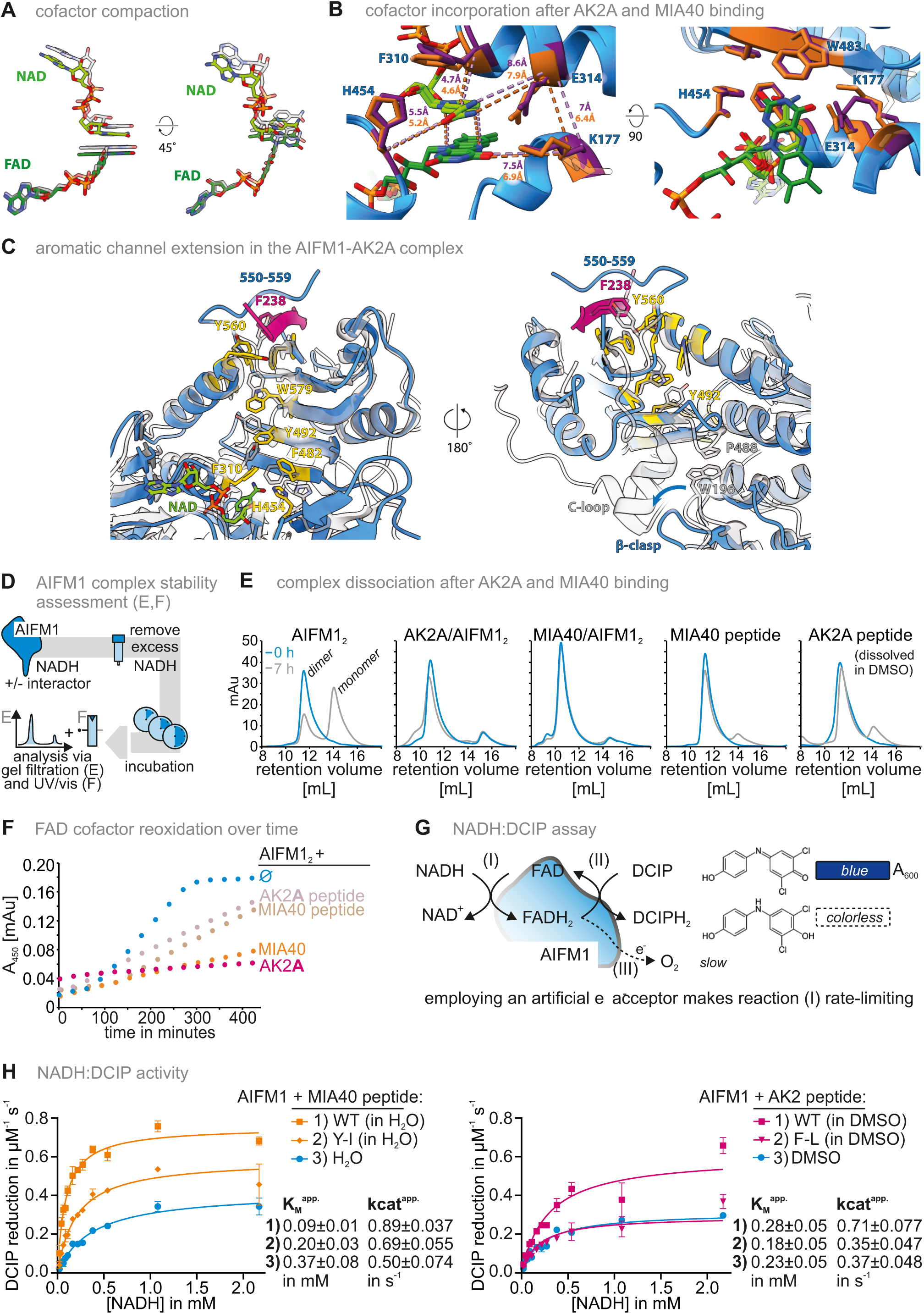
Structural impact of MIA40 and AK2A binding on AIFM1 functional domains of AIFM1 and conformations. **(A)** Isolated view of the NAD and FAD cofactor orientations in the MIA40-bound AIFM1 (light and dark green, respectively) compared to the dimer (transparent, grey overlay). **(B)** Detailed view of the AIFM1 active site. Overlay of AK2A and MIA40 bound AIFM1, residues stabilizing the cofactors shown in stick representation for AK2A (purple) and MIA40 (orange). Dashed lines: distances between the Cα atoms of the respective residues or cofactor atoms, colored according to AK2A (purple) and MIA40 (orange) bound dimer. The second observed conformation of AIFM1 K177 is visualized transparent. **(C)** Structural details of aromatic aa side chains forming the ‘aromatic tunnel’ and the conformational impact of AK2A binding (orange). Aromatic tunnel residues and the NAD binding H454 are highlighted in yellow. The AIFM1 model in the monomeric, oxidized conformation (PDB 4BV6; ^28^) is shown as a grey, transparent overlay. **(D)** Strategy for the assessment of different AIFM1 complexes *in vitro*. The AIFM1 dimer, AIFM1-MIA40, and AIFM1-AK2A complexes were established by incubating AIFM1 with AK2A, MIA40 or the respective interaction site peptides in the presence of NADH. Unbound NADH was rapidly removed using gel filtration. After different times, samples were taken and analyzed by gel filtration to visualize the respective share of complexes and AIFM1 monomer **(E)** or UV-Vis spectroscopy at 450 nm to visualize the redox state of the AIFM1 redox cofactor, FAD **(F)**. **(E)** Stability of the AIFM1 dimer, and the AIFM1-AK2A and AIFM1-MIA40 complexes. Experiment was performed as described in **(D)**. Zero and seven hours after removal of excess NADH, samples were analyzed by gel filtration and absorbance at 280 nm was used as indicator for proteins in the respective fractions. The AIFM1 dimer rapidly disassembled while complexes of AIFM1 with AK2A, MIA40 or the respective binding site peptides were strongly stabilized. **(F)** Redox state of the FAD redox cofactor. Experiment was performed as described in **(D)**. The redox state of FAD was continuously monitored over time after the removal of excess NADH. Increase of the absorbance signal indicates the oxidation of FADH_2_ over time. In the AIFM1 dimer FADH_2_ became rapidly oxidized while complexes of AIFM1 with AK2A and MIA40 or and to a lesser extent, the respective binding site peptides (AK2A peptide, MIA40 peptide) maintained FADH_2_ for longer times in the reduced state. **(G)** Strategy for the assessment of AIFM1 NADH oxidoreductase activity using 2,6 dichlorophenolindophenol (DCIP) as artificial electron acceptor. Purified AIFM1, and MIA40- or AK2A-binding site peptides are incubated in the presence of NADH to form the respective complexes. Excess NADH is removed using gel filtration. The reaction is started by mixing AIFM1 complexes with DCIP and NADH. Reduction of DCIP can be monitored by following concomitant changes in absorbance at 600 nm. **(H)** Changes in enzymatic activity of AIFM1 upon binding of MIA40- or AK2A-binding site peptides. Experiment was performed as described in **(G)**. Binding of the MIA40-binding site peptide increases the apparent k_cat_ and lowers the K_M_ towards NADH. For the AK2A-binding peptide, the apparent k_cat_ also increases while the K_M_ towards NADH remains similar. All changes are attenuated if instead of the WT peptides, peptides are used, in which single aromatic residues are mutated to aliphatic residues (MIA40-Y3I, AK2-F238L).

The NAD and FAD binding residues largely agree with the published structures of AIFM1 dimers ^26, 31^, with F310, E314 and K177 making key contacts to the cofactors. However, MIA40 binding propagates conformational changes to the active site. Specifically, the α-helix containing E314 and F310 (aa 310-326) moves closer to the cofactors by 0.2-0.6 Å (**Figure 4B, Supplementary Figure S13**). The distances between the Cα atoms of F310, E314 and H454 to NAD, and between the Cα atoms of K177, E314, and the FAD isoalloxazine are reduced in AIFM1-MIA40 as compared to the AIFM1 dimer or AIFM1-AK2A (**Figure 4B**). E314 forms hydrogen bonds with NAD and a salt bridge with K177, which then bonds with the N5 atom of the FAD isoalloxazine ring (**Supplementary Figure S13D**). Despite uncertainties in the side chain densities of K177 (probably flexible) and E314 (degraded by beam-induced damage), our AIFM1-MIA40 map suggests distinct conformations of E314 and K177 in the AIFM1 protomers of AIFM1-MIA40: one resembles the AIFM1 dimer and AIFM1-AK2A structures, while the K177 side chain of the other protomer points away from the cofactors, releasing the interactions with E314 and the FAD isoalloxazine. The lack of clearly defined side chain density in this region indicates conformational variability, reducing local resolution. In one MIA40 bound protomer, F310 and H454 show different conformations as compared to the other AIFM1 protomers solved, which may affect cofactor engagement.

Taken together, the binding of AK2A and MIA40 occurs at the same sites, but with distinct binding modes and structural consequences. AK2A binds via the addition of one β-strand and stabilizes parts of the AIFM1 C-loop. AK2A binding structurally stabilizes the AIFM1 dimer as indicated by reduced variability between AIFM1 protomers. MIA40, on the other hand, adds two β-strands and leads to a compaction of AIFM1, in particular the NAD binding domain, leading to a more compact active site including a shorter distance between the nicotinamide of NAD and the isoalloxazine of FAD. Overall, the binding of MIA40 compacts the hydrogen-bonding network around the cofactors, reducing the distance between FAD and NAD which may enhance charge transfer. When bound by MIA40, key residues involved in cofactor binding display conformational variability, with conformations varying between the AIFM1 protomers of the complex.

All our structures resemble reduced AIFM1, i.e. NAD^+^ bound after CTC formation. Consequently, we observe a key structural change upon CTC formation, which is the re-orientated conformation of aromatic side chains that form an ‘aromatic tunnel’ (F310, Y347, W351, F482, Y492, F508, Y560, and W579) connecting the central cofactor binding site and periphery of the C-terminal domain ^25^ (**Figure 4C, Supplementary Figure S14**). Alongside these conformational changes, a loop connecting two strands of the C-terminal β-sheet (aa 487-489) releases W198 of the so-called β-clasp that is part of the regulatory β-haripin (aa 190-202) (**Figure 4C, Supplementary Figure S14B**). Consequently, the C-loop is released, providing surface accessibility to the binding site of electron acceptors or a second NADH cofactor ^28^. Our structures show that both AK2A and MIA40 extend the aromatic tunnel by adding one (AK2A F238) or two (MIA40 F14, Y3) aromatic residues (**Figures 3C** and **4C, Supplementary Figure S14B)**. Through this binding mode, AK2A and MIA40 binding serves as a conformational lock, keeping the ‘aromatic tunnel’ in the NAD bound conformation and the C-loop released. Specifically, AK2A F238 and MIA40 F14 prevent re-arrangement of Y560 and, as a consequence, W579 and Y492, to arrest the loop aa 487-489 in a conformation that does not allow the stacking of P488 and W196 that contributes to β-clasp and C-loop stabilization (**Figures 3C** and **4C, Supplementary Figure S14B).**

To test the impact of these structural findings, we assessed possible changes in dimer stability and redox properties of AIFM1 induced by AK2A and MIA40 binding. The first observation was a strong impact of AK2A or MIA40 on AIFM1 dimer stability that is in line with our structural observations (**Figures 4D** and **E**). In the presence of NADH, the AIFM1 dimer forms rapidly ^15, 47^. Upon removal of excess NADH from the solution, this dimer dissolved with a half-life of about 4 hours. The binding of either purified AK2A or MIA40 proteins or AIFM1-interacting MIA40 (SYSRQEGKDRIIFVTKEDHETPSSAELVA) or AK2A (ATSKDLVMFI) peptides strongly stabilized the dimer for more than 7 hours (t_1/2_ > 7 hr, **Figure 4E**). Stabilization of the AIFM1 dimer is accompanied by inhibited re-oxidation of FADH_2_ in AIFM1 upon AK2A or MIA40 binding (**Figure 4F**). Addition of the MIA40 or AK2A peptides stabilized the reduced FAD cofactor, although to a slightly lesser extent than full-length proteins (**Figure 4F**). Another change in AIFM1 redox properties was observed in a 2,6-dichlorophenolindophenol (DCIP)-reduction assay ^25^. Without AIFM1, NADH did not reduce DCIP. Using AIFM1 alone, we observed efficient NADH oxidation with an apparent K_M_ and k_cat_ towards NADH of 0.37±0.08 mM (H_2_O, solvent for MIA40 peptides)/0.23±0.05 mM (DMSO, solvent for AK2 peptides) and 0.50±0.074 s^-1^ (H_2_O) and 0.37±0.048 s^-1^ (DMSO), respectively (**Figure 4G**). The addition of AIFM1-interacting MIA40 peptide decreased the apparent K_M_ and increased k_cat_ leading to a strong activation of AIFM1 redox activity even at physiological concentrations of NADH that are thought to be in the range of 5 – 160 µM (total NAD concentrations in the cytosol in the range of 50 – 500 µM) ^48–52^. The addition of a MIA40 peptide bearing a point mutation in an aromatic residue corresponding to Y3 of the protein (Y→I) did not result in those changes emphasizing the importance of the MIA40 interaction stabilizing the aromatic tunnel conformation. The addition of AIFM1-interacting AK2A peptides left the K_M_ mostly unchanged but increased k_cat_, implying that also in this case AIFM1 activity was increased albeit to an apparently lower extent compared to MIA40. Again, addition of an AK2A peptide with a point mutation corresponding to F238 in the protein (F→L) did not increase k_cat_. Interestingly, NADPH did not show any appreciable activity in the DCIP-reduction assay with AIFM1, and addition of AK2A or MIA40 interaction peptides did not strongly increase this negligible activity suggesting that NADH is the natural reductant employed by AIFM1, in line with previous studies (**Supplementary Figure S15B**) ^25, 28^.

Collectively, our data imply that the AIFM1 dimer is strongly stabilized by MIA40 and AK2A binding, and that the AIFM1 NADH-oxidation activity strongly increases upon MIA40 and AK2A binding. Analysis of our high-resolution cryo-EM structures provides a mechanistic explanation of how these effects are achieved. Extension of the ‘aromatic tunnel’ stabilizes the dimer-competent and C-loop released conformation that impacts both dimer stability and redox activity. Stabilization is also supported by reduced inter-protomer variability (**Supplementary Figure S11**) and the better resolved maps for AIFM1-AK2A obtained with less data than for the AIFM1 dimer, under identical sample preparation and imaging conditions (**Supplementary Figures S6-7**). AK2A further prevents the binding of the C-loop by stabilizing aa 550-558. MIA40, on the other hand, impacts the CTC and cofactor binding site, which could have an impact of the redox activity of AIFM1.

### AK2A is responsive to AIFM1 levels and complements the AK2 KO during growth on a respiratory carbon source

What is the cellular function of the AIFM1-AK2A interaction? Of the two isoforms, the AIFM1-interacting AK2A is much less abundant compared to AK2B, not only in HEK293 cells but across a panel of different cell lines (**Figures 2E** and **5A**). Moreover, its levels seem to depend on AIFM1. Deletion of AIFM1 reduces AK2A levels, while AK2B levels remain similar, and overexpression of AIFM1 increases the amounts of the AIFM1-AK2A complex (**Figures 5B** and **C**). Previous data from our group demonstrated a complex AK2 cysteine- and MIA40-dependent mechanism of AK2 import ^53^. Overall levels of AK2 seemed to be affected by loss of MIA40 ^53^ but only very mildly and not significantly affected by AIFM1 loss ^15^. A possible interpretation of these data is that import of AK2A and AK2B proceeds via the same mechanism but AK2A is less stable if it cannot establish its complex with AIFM1. Since AK2A is present in much lower amounts compared to AK2B, this might not strongly affect overall AK2 levels. The dependence of AK2A levels on AIFM1 supports that it has a function distinct from AK2B.

**Figure 5.**
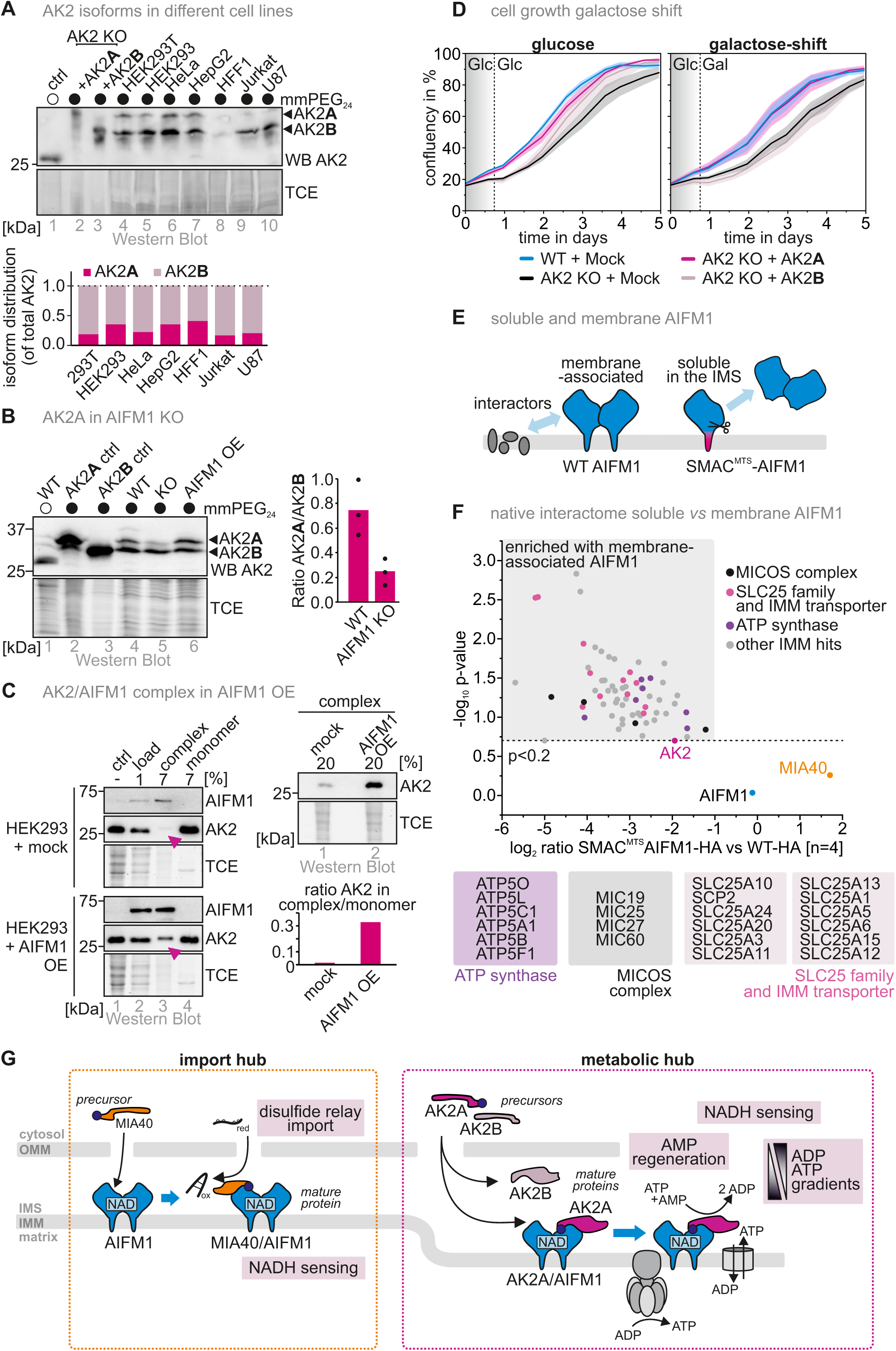
A cellular function of the AIFM1-AK2A complex in facilitating ADP/ATP exchange across the IMM. **(A)** Ratio between AK2 isoforms. As in Figure 2E, except that the thiol shift assay was performed on lysates of different cells. In all analyzed cell lines, AK2A constitutes the minor isoform with shares of 20-40%. **(B)** Levels of AK2 isoforms upon AIFM1 depletion. Experiment was performed as in **(A)** except that the experiment was performed with the indicated cell lines. Depletion of AIFM1 results in decreased AK2A levels while leaving AK2B mostly unchanged. N = 3 biological replicates. **(C)** Levels of AIFM1-AK2A complex upon AIFM1 overexpression. Experiment was performed as in Figure 2F. Overexpression of AIFM1 results in increased amounts of AIFM1-AK2A complex. **(D)** Assessment of proliferation of AK2 KO HEK293 cells complemented with either AK2A or AK2B. Cells were either grown on glucose during the entire duration of the experiment or shifted to galactose after 20 hours to test for the adaptation to the new carbon source. AK2 KO grows worse compared to WT cells on glucose as well as upon galactose shift. Complementation of the AK2 KO with AK2A rescues growth of AK2 KO cells on glucose and upon galactose shift, while AK2B rescues only glucose-grown cells. **(E)** Strategy for the assessment of AIFM1-transmembrane domain-dependent interaction partners. AIFM1 KO cell lines inducibly and stably expressing either WT AIFM1-HA or AIFM1-HA with a well-established cleavable SMAC mitochondrial targeting signal (SMAC-MTS). This latter MTS targets AIFM1 also to the IMS where it after import and processing is present in a soluble non-membrane bound state. Interactomes of these proteins will be compared **(F)** Interactome of WT AIFM1-HA and SMAC-MTS AIFM-HA. Experiment was performed as described in Figure 1B. Strikingly, many SLC25 family members and MICOS subunits as well as ATP synthase subunits were enriched on WT AIFM1. **(G)** Model. The AIFM1-AK2 interaction facilitates positioning of AK2 close to ADP/ATP transporters in the IMM. This promotes growth under conditions requiring mitochondrial respiration.

To address the physiological impact of AK2A binding to AIFM1, we next assessed the growth of AK2 KO cells complemented with either AK2A or AK2B. The AK2 KO cells were slightly impaired in growth on glucose and upon transfer of cells from glucose to galactose (galactose shift, **Figure 5D**). Complementation with AK2A and AK2B improved growth of AK2 KO on glucose albeit AK2B complementation displayed a slightly reduced growth compared to AK2A at lower cell density. This effect was more pronounced upon galactose shift, where AK2A but not AK2B fully complemented the AK2 KO.

How can the growth differences of AK2A and AK2B complemented AK2 KO cells be explained? We hypothesize that AIFM1 facilitates positioning of AK2A, providing proximity to IMM translocases. To test this, we performed interactome experiments comparing the AIFM1-HA interactome with the SMAC^MTS^-AIFM1^(103–613)^-HA interactome (**Figure 5E**). The latter AIFM1 variant is processed after import into the IMS resulting in soluble AIFM1 in the IMS, while the wildtype remains IMM bound. Interactome comparison revealed an enrichment of MICOS components, SLC25 family members and subunits of the ATP synthase with AIFM1-HA, while both variants interacted with MIA40 and AK2A (**Figure 5F**). We observed this enrichment also when we compared AIFM1-HA with mock cells (**Figures 1C** and **D**).

Collectively, we thus demonstrate that the minor AK2 isoform AK2A is sufficient to fully complement AK2 KO cells. Together with our structural analyses and the interactome studies, we speculate that bringing AK2 into proximity of ADP/ATP carriers enables an efficient transport of adenine nucleotides across the IMM which might be particularly important during shifts of carbon sources and the corresponding metabolic changes.

## DISCUSSION

### AIFM1 serves in organizing critical IMS processes

Although AIFM1 was initially associated with cell death, consistent lines of evidence demonstrate its pro-survival role as modulator of complex I biogenesis ^12^. This role in complex I assembly could be explained through a series of elegant studies establishing a link between AIFM1 and the mitochondrial disulfide relay ^13–15, 22, 23^. AIFM1 facilitates the import and binding of MIA40 in the IMS. This interaction enables the efficient import of specific disulfide relay substrates, including NDUFS5, which is part of complex I modules and its insertion has been identified as an AIFM1-dependent rate-limiting step in complex I assembly ^15^. Our cryo-EM structure confirmed that MIA40 interacts with AIFM1 via its N-terminal stretch of aa 2-20 and suggests that this is the main site of interactions between AIFM1 and MIA40 (**Figures 3B** and **C**).

In addition to MIA40, we identified previously unknown, high-confidence interaction partners of AIFM1: AK2 and subunits of the MICOS complex (**Figure 1**). The interaction with MICOS components might either indicate positioning of AIFM1 by the MICOS complex or a role of AIFM1 in MICOS assembly and function, which will be exciting to investigate in future studies. Here, we thoroughly investigated the interaction between AIFM1 and AK2, and demonstrated that only isoform AK2A is able to interact with AIFM1. AK2A is less abundant compared to AK2B, and we found that it is stabilized by AIFM1 binding (**Figure 5B**). Interestingly, AK2A and MIA40 bind the same site in the AIFM1 C-terminal domain by β-strand complementation. However, an N-terminal stretch of MIA40 binds AIFM1, whereas AK2A uses its last seven amino acids, which are absent in the isoform AK2B. Interestingly, a human patient in which K233 was mutated to a termination codon, essentially resulting in expression of only AK2B, suffered from reticular dysgenesis emphasizing the physiological importance of the AIFM1 AK2A interaction ^45^. The dissociation constant of the AIFM1-AK2A interaction is by a factor of three higher than the K_d_ of AIFM1-MIA40, suggesting that MIA40 outcompetes AK2A for AIFM1 binding. We confirmed this conclusion by our gel filtration assays using purified proteins as well as cell lysates. In contrast to AK2B, which appears to be imported and maintained in the IMS independent of AIFM1^15^, AK2A depends on the interaction with AIFM to be stably present in the IMS. The competition with MIA40 for AIFM1 binding might therefore indicate that in conditions of limiting amounts of dimeric AIFM1, a considerable amount of AK2A might remain unbound to AIFM1, become destabilized as a consequence potentially resulting in changes of metabolic signaling from mitochondria.

### AIFM1 forms complexes with AK2A and MIA40 of varying stoichiometry

We determined a stoichiometry of one MIA40 or AK2A per AIFM1 dimer by ITC (**Figure 2H**), while gel filtration after long-term co-incubation indicated that tetramers of two MIA40 or AK2A protomers per AIFM1 might also be possible. Accordingly, our cryo-EM reconstructions and published structural data ^31^ show two molecules of AK2A or MIA40 bound to the AIFM1 dimer. The inter-protomer variability of AIFM1 that we observe within the AIFM1 dimer and AIFM1-MIA40 suggests a degree of structural asymmetry within the complex. This is also reflected in the differences in local cryo-EM densities of the sixth β-strand of AIFM1 added to the C-terminal β-sheet in the absence of interactors, as well as in the protomer differences observed by x-ray crystallography ^25, 31^. The inherent structural variability within the AIFM1 dimer may be linked to the variability within the AIFM1-MIA40 complex that we observe in our cryo-EM structure, which could be induced by MIA40 initially binding to one of the AIFM1 protomers. The possibility of one or two MIA40 or AK2A molecules binding to an AIFM1 dimer also leaves open the possibility of a mixed complex comprising AIFM1 and both MIA40 and AK2A.

### MIA40 and AK2A affect the redox and NADH turnover properties of AIFM1

Besides the relevance of the AIFM1-AK2A interaction for adenylate kinase function, our results also include implications for the oxidoreductase function of AIFM1. NAD(P)H dependent AIFM1 dimerization is a prerequisite for its interaction with either MIA40 ^13, 15, 47^ or AK2A (**Figure 2G**). Over time, NAD(P)H becomes slowly oxidized resulting in the dissociation of the AIFM1 dimer. Our structural data provide an explanation of how AK2A and MIA40 stabilize AIFM1 dimers. By binding to the C-terminal domain of AIFM1, MIA40 and AK2A cap the aromatic tunnel by adding two and one aromatic side chains, respectively. We propose that this binding event stabilizes the conformation of these aromatic side chains that is a structural hallmark of the reduced, dimeric form of AIFM1. Furthermore, a critical structural element, the aa 487-490 loop is kept in a conformation that prevents closing of the β-clasp, i.e. a conformation of the β-hairpin (aa 190-202) that regulates the stabilization of the C-loop. AK2A further interacts with aa 550-558 and thereby reduces the ability of the C-loop to engage with its inhibitory binding site the AIFM1 C-terminal domain. Thereby, MIA40 and AK2A binding favors a detached state of the C-loop, keeping the electron acceptor binding site solvent exposed.

AIFM1 has been shown to act as NADH oxidase or NADH:ubiquinone oxidoreductase which enables it to feed electrons into the respiratory chain. However, AIFM1 is normally not very efficient as NADH oxidizing enzyme as it has relatively high K_M_ values for NADH in the DCIP assay (K_M_ = 370 µM) well above the physiological levels of the metabolite (5 – 160 µM NADH ^48–52^). Using an artificial, commonly employed electron acceptor, we demonstrated that MIA40 or AK2A binding significantly increased NADH oxidation activity in the physiological NADH range. How this translates to the potential physiological electron acceptor ubiquinone remains to be shown, but it might imply that AIFM1 requires AK2A or MIA40 binding to efficiently sustain the electron flow from cytosolic NADH into the respiratory chain to promote ATP biosynthesis. The increased oxidoreductase activity of the complex can most likely be explained by the impact of AK2A and MIA40 on the stability of the dimer and detached/disordered conformation of the C-loop, thus favoring the surface accessible conformation of the electron acceptor binding site. AK2A additionally binds part of the C-loop, further preventing its (re-) attachment. MIA40, on the other hand, may impact the oxidoreductase activity of AIFM1 by other means, namely by propagating a compaction to the CTC, reducing the FAD-NAD distance. Therefore, MIA40 and AK2A have a dual impact on AIFM1 function, affecting dimer stability and oxidoreductase activity, by directly impacting the concerted structural rearrangements that mediate dimerization and C-loop release upon CTC formation.

### AIFM1-AK2A complex formation positions AK2A close to ADP/ATP translocases

Collectively, our data are in line with a model (**Figure 5G**) in which the presence of NADH would allow AIFM1 dimer formation and subsequent AK2A binding. Growth experiments on different carbon sources revealed that the ability of AK2A to interact with AIFM1 is crucial for its cellular function. In contrast to AK2A, the expression of AK2B which lacks the C-terminal residues required for AIFM1 interaction did not allow to rescue the growth defect of an AK2 deletion, although both isoforms showed similar enzymatic profiles (**Supplementary Figure S3**). In light of our membrane domain-dependent AIFM1 interactome, which revealed many SLC25 members including ADP/ATP transporters and the ATP synthase, we propose that the AIFM1-AK2A interaction is critical for positioning AK2A close to a metabolic hub of ADP/ATP translocases and the ATP synthase to facilitate efficient ADP-ATP exchange across the IMM and the IMS. Since the carriers serve as reversible antiporters for ADP and ATP, AK2A close by would be important two-fold: firstly, to convert any remaining AMP, generated in high amounts during cellular processes such as fatty acid activation or aminoacyl tRNA synthesis, into ADP using the high amounts of ATP in proximity to the carrier. This is supported by the low K_M_ of AK2 towards AMP ^53^ and avoids accumulation of an adenine nucleotide that cannot be transported across the IMM. Secondly, AK2 in close proximity to the carriers contributes to maintain through its phosphotransferase activity a steady adenine nucleotide gradient ^33, 34, 37^, rendering ADP/ATP exchange by the respective carriers more efficient.

How do fluctuations in NADH/NAD^+^ ratios, as they occur e.g. during starvation or varying glycolysis activity, influence the activity of the AIFM1-AK2A complex? Since both AIFM1-AK2A and AIFM1-MIA40-complexes are rather stable in the absence of NADH (**Figures 4D** and **F**), we assume that short-term fluctuations leave both complexes intact. Interestingly, AK2 is stringently regulated by AMP inhibition in a range that might become relevant under starvation conditions ^53, 54^. Thus, during short-term starvation and upon increasing AMP levels, AK2A activity in the AIFM1-AK2A complex might be switched off decreasing the efficiency of ADP-ATP exchange across the IMM and the IMS. Lower levels of NADH during a longer time period might then also affect the initial formation of the AIFM1-AK2A complex.

In summary, we demonstrate that AIFM1 contributes to the organization of the IMS. Importantly, AIFM1 regulates the two branches of the OXPHOS system, since it controls Complex I biogenesis and ADP availability that is key to maintain the activity of the ATP synthase.

## MATERIALS AND METHODS

### Plasmids, cell lines and chemical treatments of cells

For plasmids, cell lines, antibodies and further tools used in this study, see **Tables S1-S5**. Cells were cultured in DMEM supplemented with 8% fetal calf serum (FCS) at 37°C under 5% CO_2_. For cycloheximide (CHX) chase experiments, cells were treated with 100 µg/mL CHX dissolved in DMSO. For the generation of stable, inducible HEK293 cell lines, the Flp-In T-REx-293 cell line was used with the Flp-In T-REx system (Invitrogen). For generation of stable, inducible HEK293T cell lines, the PiggyBac Transposon system (System Biosciences, BioCat) was used.

Expression of constructs was induced using 1 µg/mL doxycycline (Flp-In T-Rex) or 30 µg/mL cumate (PiggyBac Transposon) for the indicated time points. Expression of SMAC^MTS^-AIFM1-HA in HEK293T cells was induced with only 15 µg/mL to obtain comparable protein levels.

### Generation of HEK293 knockout cells

HEK293 knockout cell lines were generated using the pSpCas9(BB)-2A-GFP (PX458) CRISPR/Cas9 construct (a gift from F. Zhang; Addgene, plasmid 4813; ^55^) as described previously ^56^. In brief, CRISPR/Cas9 gRNAs were designed for gene disruption using CHOPCHOP software ^57^. Transfections were performed using Lipofectamine LTX (Thermo Fisher Scientific) and green fluorescent cells were individually sorted.

### Immunoprecipitation

Immunoprecipitations were carried out under native lysis conditions. The cells were washed with PBS, supplemented with 20 mM NEM (N-Ethylmaleimide). After incubation in PBS supplemented with 20 mM NEM for 15 minutes, the cells were mechanically detached by scraping and sedimented with 800 x g for 5 minutes at 4°C. The cells were gently lysed in ice-cold native IP lysis buffer (100 mM sodium phosphate pH 8.0, 100 mM sodium chloride, 1% (v/v) Triton X-100, 0.2 mM PMSF) for 1 h on ice. Lysate was cleared by centrifugation 22,000 x g for 1 h at 4°C. Supernatant was transferred to a prewashed agarose matrix and incubated for 3.5 to16 h at 4°C on a tumbling shaker. Afterwards, beads were triply washed with the IP lysis buffer, containing Triton X-100 and once finally washed with IP lysis buffer without Triton X-100. Precipitated proteins were eluted from the agarose matrix by addition of Laemmli buffer (2% SDS, 60 mM Tris-HCl pH 6.8, 10% glycerol, 0.0025% bromophenol blue) and heating up to 96°C for 2 times 4 min.

### SILAC-based mass spectrometry

The experiment was performed as described in ^22^. Cells were subcultured and passaged in SILAC-DMEM (Thermo Fisher), supplemented with 10% dialysed FBS (Gibco, Invitrogen), 1% L-glutamine (PAN Biotech), containing either L-arginine or L-arginine-13C6-15N4 (42 mg/L), and L-lysine or L-lysine-13C615N2 (73 mg/L), and 27.3 mg/L proline. After immunoprecipitation, samples were eluted in SDS-PAGE loading buffer containing 1 mM DTT (Sigma-Aldrich) and alkylated using 5.5 mM iodoacetamide (Sigma-Aldrich). Protein mixtures were separated by SDS-PAGE, gel lanes were cut into 10 equal slices, proteins therein were in-gel digested with trypsin (Promega) and the resulting peptide mixtures were processed on STAGE tipps. Mass spectrometric measurements were performed on an LTQ Orbitrap XL mass spectrometer (Thermo Fisher Scientific) coupled to an Agilent 1200 nanoflow-HPLC (Agilent Technologies GmbH) as described ^58^. The MS raw data files were uploaded into the MaxQuant software ^59^. A full length IPI human database containing common contaminants such as keratins and enzymes used for in-gel digestion was employed. Methionine oxidation, protein amino-terminal acetylation, carbamidomethyl cystein and NEM cysteine were set as variable modifications. Double SILAC was chosen as quantitation mode. The MS/MS tolerance was set to 0.5 Da. Peptide lists were further used by MaxQuant to identify and relatively quantify proteins using the following parameters: peptide, and protein false discovery rates (FDR) were set to 0.01, maximum peptide posterior error probability (PEP) was set to 0.1, minimum peptide length was set to 6, minimum number peptides for identification and quantitation of proteins was set to one which must be unique, and identified proteins have been re-quantified.

### In vitro protein-protein profiling of putative AIFM1 interactors by using microchips

To find novel interactors of AIFM1, a high-content protein microarray was performed using ProtoArray™ Human Protein Microarrays v5.1 (Thermo Fisher Scientific) containing ∼ 9.000 N-terminal Glutathione S-Transferase (GST)-tagged human proteins extracted from transfected insect cells. As described in one of our prior studies ^60^, each ProtoArray™ plate was placed at 4°C for equilibration for at least 15 minutes prior to blocking. Plates were then blocked using 5 mL blocking solution (50 mM HEPES, 200 mM NaCl, 0.08% Triton X-100, 25% glycerol, 20 mM glutathione, 1.0 mM DTT, 1X Synthetic Block) at 4 °C for 1 h on a shaker at 50 rpm. After incubation, the blocking solution was aspirated and plates were incubated with recombinant AIFM1 protein (concentration of 5 ng/mL and 50 ng/mL) diluted in probe buffer (1X PBS, 0.1% Tween-20, 1X Synthetic Block), while one microarray (negative control) was exposed only to probe buffer for 90 min at 4 °C. Afterwards, microplates were washed 5 times for 5 min with wash buffer (1X PBS, 1X Synthetic Block, 0.1% Tween 20). After washing, microplates were incubated with primary antibody in probe buffer for 90 min at 4 °C, washed 5 times in probe buffer and incubated with Alexa Fluor™ 647-conjugated goat anti-rabbit IgG (Thermo Fisher Scientific, #A21244: Lot 1654324, 1 μg/mL in probe buffer) for 90 min at 4 °C. Plates were then washed 5 times 5 min with wash buffer. To remove the residual salt, each plate was quickly washed with distilled water and dried by centrifuging at 200 g for 1 min. ProtoArray™ plates were scanned using an Axon 4000B fluorescent microarray scanner (Molecular Devices). Hits were considered based on the following criteria: (a) the fluorescent intensity value of the hits should be at least 20-fold higher than the corresponding negative control; (b) the normalized fluorescent signal was greater than 3 standard deviations; (c) the signal-to-noise ratio was higher than 0.5 and (d) the replicate spot coefficient of variation (CV) was lower than 65%.

### Immunoblotting and image acquisition

Samples were prepared in Laemmli buffer containing 50 mM dithiothreitol (DTT), and heat denatured for 5 min at 96 °C and DNA degraded by sonification (50% amplitude, 16 cycles). Protein samples were analyzed by SDS-PAGE and immunoblotting. The addition of 2,2,2-trichloroethanol (TCE) to the SDS-PAGE gel allowed for visualization of proteins and as a loading control. The immunoblotting images were detected using the ChemiDoc Touch Imaging system (Bio-Rad).

### In vitro AK2 activity assay

The activity assay was carried out by coupling the AK2 reaction to hexokinase (HK) and glucose-6-phosphate dehydrogenase (G6PDH, HK/G6PDH mix from Roche). In this assay, AK2 provides the ATP for glucose phosphorylation by HK, followed by NADP^+^ reduction to NADPH and an increase in absorbance at 340 nm. The reaction conditions were as follows: 58 mM glycylglycine pH 7.4, 10 mM MgCl_2_, 0.006% BSA, 0.25 mM NADP^+^, 20 mM glucose. The concentration of AK2 was set to 8 nM. All measurements were performed in triplicates in 96-well plates and read in a CLARIOstar microplate reader set to 25°C. A measurement without AK2 was performed simultaneously to all measurements to allow subtraction of the background reaction.

### Analytical size-exclusion chromatography

Analytical size-exclusion chromatography was performed under native conditions to examine protein complexes between intact proteins. Cells were washed with 1x PBS and mechanically detached by scraping. Cells were sedimented at 500 g for 5 min. Pellets were resolved in 660 μL native lysis buffer (100 mM sodium phosphate pH 8.0, 100 mM sodium chloride, 1% (v/v) Triton X-100), supplemented with 0.2 mM PMSF. Cells were lysed for 1 h on ice, and the lysate was cleared by centrifugation. Lysate was loaded on a HiLoad™ 16/600 Superdex 200 preparation grade gel filtration column and installed in a liquid chromatography system (Aekta Purifier) from GE Healthcare. A protein size standard was used as a reference, covering a range from 1.35 kDa to 670 kDa (#1511901, Bio-Rad).

### Assay to address thiol redox states of proteins

The redox state assay was performed as previously described ^61^. Cells were lysed in Laemmli buffer and oxidized cysteines were reduced by the addition of 10 mM TCEP (Tris(2-carboxyethyl)phosphine) and incubated at 96°C for 10 min. Following, the newly reduced cysteines were modified with the alkylating agent mmPEG24. Alkylation with 15 mM mmPEG24 was carried out for 1 h at room temperature. Subsequently, samples were separated by SDS–PAGE and analyzed by western blot, followed by immunoblotting.

### Peptide synthesis

The peptides SYSRQEGKDRIIFVTKEDHETPSSAELVA-NH2 (MW_calc_ = 3292.58 Da; MW_exp_ = 3293.20, final purity 91 %), SICRQEGKDRIILVTKEDHETPSSAELVA-NH2 (MW_calc_ = 3224.61 Da; MW_exp_ = 3325.74, final purity 78 %), ATSKDLVMFI-NH2 (MW_calc_ = 1123.36, MW_exp_ = 1024.09 final purity 98 %) and ATSKDLVMLI-NH2 (MW_calc_ = 1090.35, MW_exp_ = 1090.08, final purity 98 %) were synthesized by solid-phase peptide synthesis on a peptide synthesizer (Syro I, MultiSynTech) using the fluorenylmethoxycarbonyl (Fmoc)/*tert*-butyl strategy on a Rink amide resin (0.48 mmol/g, 15 μmol scale). The amino acid coupling steps were performed twice for each amino acid using eight equivalents each of OxymaPure (2-cyano-2(hydroxyamino)acetate), DIC (dicyclohexylcarbodiimide), and the respective Fmoc-protected amino acid in DMF (dimethylformamide). The protecting group was removed by incubating the resin first in 40% piperidine in DMF followed by 20% piperidine in DMF. Peptides were cleaved from the resin using a mixture of trifluoroacetic acid/ thioanisole/ 1, 2-ethanedithiol (90:7:3, v/v/v). The crude peptides were purified by reverse-phase HPLC using a linear gradient of 10–60% B in A (A: water/0.1 % TFA; B: acetonitrile (ACN)/0.1 % TFA) over 45 min.

### Protein purification

Recombinant proteins were expressed from the indicated plasmids (**Table S2**) in Rosetta2 *E. coli* strains. Bacterial growth was conducted in LB media (for AIFM1 supplemented with riboflavin and FAD) shaking at 37°C and 180 rpm. AIFM1(103-613) expression was induced with 1.0 mM IPTG and incubated for further 16 h before harvesting. AK2A C40,232S and AK2B C40S expression was induced with 0.1 mM IPTG and incubated further 16 h at 25°C. MIA40 C4S,C53S,C55S expression was induced with 0.5 mM and incubated for further 3 h. Cells were harvested on ice in PBS and stored at −20°C. The 6xHis-tagged constructs were purified by Immobilized Metal Affinity Chromatography using Ni Sepharose (6 Fast Flow, GE). The bacterial lysate was bound to beads in binding buffer supplemented with 10 mM imidazole at 4°C. Beads were washed with binding buffer supplemented with 20 mM imidazole prior to elution with 150 mM imidazole. Imidazole was removed using PD-10 columns (Cytiva) and the proteins stored at 4 °C.

### In vitro reconstitution of the AIFM1-MIA40 and AIFM1-AK2A interactions

The AIFM1 AK2A or MIA40 complex, respectively, was reconstituted by combining the recombinant proteins in a 1:2 to 1:4 (AIFM1:AK2A/MIA40) molar ratio in 100 mM NaCl, 20 mM Tris/CL pH = 7.4 in presence of 0.1 mM NADH. After incubation for 20 minutes and centrifugation, the respective complex was separated from monomeric proteins on a 16/600 Superdex 200 PG or on a Superdex 200 Increase 10/300 GL column.. All steps were performed at 4°C.

### Isothermal titration calorimetry (ITC)

Isothermal titration calorimetry was performed at 25°C on a MicroCal Auto-ITC200 (Malvern, United Kingdom). For analysis of the interaction between AIFM1 and peptides, 6xHis-tagged AIFM1(103–613) was dialyzed against PBS pH 7.4 supplemented with 0.1 mM NADH at 4°C for 17 h. Lyophilized peptide was dissolved in dialysis buffer to a final concentration of 250 - 300 μM. Ligand proteins were dialyzed in the same batch of buffer as AIFM1(103-613) als used in a concentration of 250 – 300 µM. The concentration of receptor in the sample cell was 30 μM. Measurements were carried out by 2 μl injections of the peptide into the cell with an injection duration of 4 s. Ultimately, 19 injections were performed during the titration.

### Cryo-EM grid preparation and data collection

Before cryo-EM, sample quality was assessed by negative staining electron microscopy as previously described ^62^ (data not shown). For cryo-grid preparation, 3 µL of purified protein was applied to an UltrAuFoil® R 1.2/1.3 grid (Quantifoil) that had been glow-discharged for 1 minute and 45 seconds. The grids were then blotted for 4 seconds at 100% humidity and 8°C, followed by plunging in liquid ethane cooled by liquid nitrogen using a Vitrobot Mark IV (Thermo Fisher Scientific). The prepared grids were stored in liquid nitrogen until use.

Cryo-EM data for AIFM1-MIA40 were collected using a Titan Krios G3i (Thermo Fisher Scientific), while data for AIFM1 dimer and AIFM1 with AK2 were collected using Titan Krios G4 (Thermo Fisher Scientific), all operated at 300 kV with a 35° tilt to overcome preferred orientation using EPU (Thermo Fisher Scientific).

Two datasets were collected for AIFM1-MIA40: 1907 raw movies for the first dataset and 2084 for the second, using a Falcon III direct electron detector with a pixel size of 0.654 Å/pixel. The total electron dose was 50.82 e/Å² for the first dataset and 50.57 e/Å² for the second, distributed over 48 frames with a defocus range of −0.6 to −2.6 µm.

For the AIFM1 dimer, 15,440 raw movies with a pixel size of 0.46 Å/pixel and for AIFM1 with AK2A, 4542 movies with a pixel size of 0.58 Å/pixel were collected using a Falcon 4i equipped with a Selectris energy filter (Thermo Fisher Scientific). These movies were stored in electron-event representation (EER) format, with a total dose of 50 e/Å² distributed over 468 frames and a defocus range of −0.7 to −1.7 µm.

### Cryo-EM data processing

All datasets were processed using cryoSPARC (V4.4) ^63^. The workflows are described in detail in **Supplementary Figures S6-8**. In summary, the movies were pre-processed with patch-based motion correction and CTF estimation. Initial blob picking and 2D classification were performed to identify classes with distinguishable features, which were then used to train the TOPAZ picker ^64^. After further 2D classification, particles were divided into high and low-defocus groups for further TOPAZ training. Multiple iterations of TOPAZ picking, 2D classification and ab initio reconstruction were performed, followed by homogeneous or non-uniform refinement of good reconstructions.

For the AIFM1 dimer, two initial models containing 149,783 and 188,760 particles, respectively, were pooled, classified using ab initio reconstruction, and refined using non-uniform refinement. The initial models of the AIFM1-AK2A complex (229,483 particles) and the AIFM1-MIA40 complex (310,098 particles), the particles were subjected to another round of TOPAZ training, 2D classification, and ab initio reconstruction, followed by either non-uniform or homogeneous refinement.

Particles of AIFM1 with MIA40 were extracted with a box size of 420 pixels, while the AIFM1 dimer and AIFM1 with AK2A were extracted with a box size of 416 pixels. All refinements were eventually processed with Reference Motion correction resulting in resolutions of 2.8 Å for the AIFM1 dimer (277,866 particles), 2.4 Å for AIFM1 with MIA40 (291,656 particles) and 2.6 Å for AIFM1 with AK2A (307,496 particles). Statistics on data collection and validation are given in **Supplementary Figures S6-8**.

### Model building and refinement

For all AIFM1 structures obtained in this study, an AlphaFold2 model was used to obtain an initial model and then crossreferenced with previously published human and mouse AIFM1 models (PDB:4BUR and PDB:3GD4). Residues were adjusted in Coot ^65^. and models were iteratively refined and adjusted using PHENIX (V1.21) ^66^ and Coot. For better visualization during initial model building, maps were processed with DeepEMhancer ^67^. Maps and models were visualised using ChimeraX ^68^. Statistics on data collection and validation reports were automatically generated using MolProbity within Phenix ^69^.

### NAD(P)H: 2,6 dichlorophenolindophenol (DCIP) activity assay

AIFM1 catalyzes the efficient reduction of DCIP. In a two-step reaction, NADH first reduces the FAD cofactor in AIFM1, from which subsequently electrons are transferred onto DCIP. Using DCIP as electron acceptor, the latter step is faster than the first one allowing to observe the AIFM1-dependent oxidation of NADH in dependence of differing NADH concentrations. The enzymatic activity of AIFM1 as NAD(P)H:DCIP oxidoreductase was measured in 20 mM NaCl, 20 mM Tris/Cl pH = 7.4 at 25°C. Recombinant AIFM1 was used in a final concentration of 800 – 927 nM, The concentration of DCIP was kept constant at 200 µM and the indicated peptides at a final concentration of 3.3 µM. The reaction was started by adding NAD(P)H and the DCIP reduction monitored at 600 nm in a plate reader (CLARIOstar, BMG Labtech). The background reaction in absence of AIFM1 for each NAD(P)H concentration was subtracted from the data.

### Quantification and statistical analysis

Intensity of immunoblot signals were quantified using Image Lab (Biorad). Error bars in figures represent standard deviation. The number of experiments is reported in the figure legend.

## Supporting information

Supplemental Information complete

Structure table

## DATA AVAILABILITY

The datasets generated during and/or analysed during the current study are available from the corresponding author on reasonable request. The mass spectrometry proteomics data have been deposited to the ProteomeXchange Consortium via the PRIDE ^70^ partner repository with the dataset identifier PXD055617.

## ACKNOWLEDGEMENTS

The Deutsche Forschungsgemeinschaft (DFG, German Research Foundation) funds research in the Laboratory of JR through the grants RI2150/5-1 project number 435235019, SPP2453 project number 541742459, RTG2550/1 project number 411422114, CRC1218 - project number 269925409, and CRC1678 - project number 520471345. SP is funded by CMMC core funding (JRG XI), by the DFG - 455 SFB1430 - Project-ID 424228829 and the CANTAR network funded by the Ministry of Culture and Science of the state of Northrhine-Westphalia. DB is supported by DZNE core budget and the SPP2453 project number 541742459. DB is a member of the ETERNITY project consortium, funded by the European Union through Horizon Europe Marie Skłodowska-Curie Actions Doctoral Networks (MSCA-DN) under the grant number 101072759. We thank the CECAD Proteomics Facility for the analysis of proteome data; the facility is supported by the large instrument grant INST 216/1163-1 FUGG by the DFG. We thank the CECAD Imaging as well as the StruBiTEM cryo-EM facilities, in particular Monika Gunkel and Elmar Behrmann for their support. Cryo-EM data of the AIFM1 dimer and AIFM1-AK2A were collected at the cryoEM center of the University of Münster (cryoEM SoN), which is funded by the DFG – Projektnummer 496113311. We thank Christos Gatsogiannis and Alexander Neuhaus for support. We thank Ms Christiane Bartling-Kirsch (DZNE) and Dr. Lena Wischhof (DZNE) for their important help with this study. We thank Kathrin Ulrich, Matthias Weith and members of the Riemer lab for critical reading of the manuscript. We thank Anja Wittmann and Anika Seiler for technical support throughout the project.

## AUTHOR CONTRIBUTIONS

JR designed the study. RAR designed and cloned constructs. RAR, KL designed and generated cell lines. RAR, DS, SM, PG, KW, SG carried out the biochemical experiments. JuR performed immunoprecipitations and label free proteomics experiments. CP, JD performed SILAC proteomics experiments. SP, EP solved the cryo-EM structures. SG performed cell proliferation assays. DE, MM and DB performed *in vitro* protein-protein profiling experiments and *C. elegans* experiments. KS, IN designed and synthesized peptides. All authors analysed data. JR, SP wrote the first manuscript draft. All authors provided input to the manuscript and proofread it.

## REFERENCES

1. Herrmann, J.M. & Riemer, J. Apoptosis inducing factor and mitochondrial NADH dehydrogenases: redox-controlled gear boxes to switch between mitochondrial biogenesis and cell death. Biol Chem 402, 289–297 (2021).

2. Novo, N., Ferreira, P. & Medina, M. The apoptosis-inducing factor family: Moonlighting proteins in the crosstalk between mitochondria and nuclei. IUBMB Life 73, 568–581 (2021).

3. Reinhardt, C. et al. AIF meets the CHCHD4/Mia40-dependent mitochondrial import pathway. Biochim Biophys Acta Mol Basis Dis 1866, 165746 (2020).

4. Wischhof, L., Scifo, E., Ehninger, D. & Bano, D. AIFM1 beyond cell death: An overview of this OXPHOS-inducing factor in mitochondrial diseases. EBioMedicine 83, 104231 (2022).

5. Bano, D. & Prehn, J.H.M. Apoptosis-Inducing Factor (AIF) in Physiology and Disease: The Tale of a Repented Natural Born Killer. EBioMedicine 30, 29–37 (2018).

6. Klein, J.A. et al. The harlequin mouse mutation downregulates apoptosis-inducing factor. Nature 419, 367–374 (2002).

7. van Empel, V.P. et al. Downregulation of apoptosis-inducing factor in harlequin mutant mice sensitizes the myocardium to oxidative stress-related cell death and pressure overload-induced decompensation. Circ Res 96, e92–e101 (2005).

8. van Empel, V.P. et al. EUK-8, a superoxide dismutase and catalase mimetic, reduces cardiac oxidative stress and ameliorates pressure overload-induced heart failure in the harlequin mouse mutant. J Am Coll Cardiol 48, 824–832 (2006).

9. Ghezzi, D. et al. Severe X-linked mitochondrial encephalomyopathy associated with a mutation in apoptosis-inducing factor. Am J Hum Genet 86, 639–649 (2010).

10. Bertan, F. et al. Comparative analysis of CI- and CIV-containing respiratory supercomplexes at single-cell resolution. Cell Rep Methods 1, 100002 (2021).

11. Wischhof, L. et al. A disease-associated Aifm1 variant induces severe myopathy in knockin mice. Mol Metab 13, 10–23 (2018).

12. Vahsen, N. et al. AIF deficiency compromises oxidative phosphorylation. EMBO J 23, 4679–4689 (2004).

13. Hangen, E. et al. Interaction between AIF and CHCHD4 Regulates Respiratory Chain Biogenesis. Mol Cell 58, 1001–1014 (2015).

14. Meyer, K. et al. Loss of apoptosis-inducing factor critically affects MIA40 function. Cell Death Dis 6, e1814 (2015).

15. Salscheider, S.L. et al. AIFM1 is a component of the mitochondrial disulfide relay that drives complex I assembly through efficient import of NDUFS5. EMBO J 41, e110784 (2022).

16. Zarges, C. & Riemer, J. Oxidative protein folding in the intermembrane space of human mitochondria. FEBS Open Bio (2024).

17. Geldon, S., Fernandez-Vizarra, E. & Tokatlidis, K. Redox-Mediated Regulation of Mitochondrial Biogenesis, Dynamics, and Respiratory Chain Assembly in Yeast and Human Cells. Front Cell Dev Biol 9, 720656 (2021).

18. Al-Habib, H. & Ashcroft, M. CHCHD4 (MIA40) and the mitochondrial disulfide relay system. Biochem Soc Trans 49, 17–27 (2021).

19. Backes, S. & Herrmann, J.M. Protein Translocation into the Intermembrane Space and Matrix of Mitochondria: Mechanisms and Driving Forces. Front Mol Biosci 4, 83 (2017).

20. Modjtahedi, N., Tokatlidis, K., Dessen, P. & Kroemer, G. Mitochondrial Proteins Containing Coiled-Coil-Helix-Coiled-Coil-Helix (CHCH) Domains in Health and Disease. Trends Biochem Sci 41, 245–260 (2016).

21. Stojanovski, D., Bragoszewski, P. & Chacinska, A. The MIA pathway: a tight bond between protein transport and oxidative folding in mitochondria. Biochim Biophys Acta 1823, 1142–1150 (2012).

22. Petrungaro, C. et al. The Ca(2+)-Dependent Release of the Mia40-Induced MICU1-MICU2 Dimer from MCU Regulates Mitochondrial Ca(2+) Uptake. Cell Metab 22, 721–733 (2015).

23. Murschall, L.M. et al. The C-terminal region of the oxidoreductase MIA40 stabilizes its cytosolic precursor during mitochondrial import. BMC Biol 18, 96 (2020).

24. Mate, M.J. et al. The crystal structure of the mouse apoptosis-inducing factor AIF. Nat Struct Biol 9, 442–446 (2002).

25. Churbanova, I.Y. & Sevrioukova, I.F. Redox-dependent changes in molecular properties of mitochondrial apoptosis-inducing factor. J Biol Chem 283, 5622–5631 (2008).

26. Sevrioukova, I.F. Redox-linked conformational dynamics in apoptosis-inducing factor. J Mol Biol 390, 924–938 (2009).

27. Sorrentino, L. et al. Key Role of the Adenylate Moiety and Integrity of the Adenylate-Binding Site for the NAD(+)/H Binding to Mitochondrial Apoptosis-Inducing Factor. Biochemistry 54, 6996–7009 (2015).

28. Ferreira, P. et al. Structural insights into the coenzyme mediated monomer-dimer transition of the pro-apoptotic apoptosis inducing factor. Biochemistry 53, 4204–4215 (2014).

29. Brosey, C.A. et al. Defining NADH-Driven Allostery Regulating Apoptosis-Inducing Factor. Structure 24, 2067–2079 (2016).

30. Villanueva, R. et al. Key Residues Regulating the Reductase Activity of the Human Mitochondrial Apoptosis Inducing Factor. Biochemistry 54, 5175–5184 (2015).

31. Fagnani, E. et al. CHCHD4 binding affects the active site of apoptosis inducing factor (AIF): Structural determinants for allosteric regulation. Structure 32, 594–602 e594 (2024).

32. Kettwig, M. et al. From ventriculomegaly to severe muscular atrophy: expansion of the clinical spectrum related to mutations in AIFM1. Mitochondrion 21, 12–18 (2015).

33. Dzeja, P. & Terzic, A. Adenylate kinase and AMP signaling networks: metabolic monitoring, signal communication and body energy sensing. Int J Mol Sci 10, 1729–1772 (2009).

34. Dzeja, P.P., Zeleznikar, R.J. & Goldberg, N.D. Adenylate kinase: kinetic behavior in intact cells indicates it is integral to multiple cellular processes. Mol Cell Biochem 184, 169–182 (1998).

35. Noma, T. Dynamics of nucleotide metabolism as a supporter of life phenomena. J Med Invest 52, 127–136 (2005).

36. Ruprecht, J.J. & Kunji, E.R.S. The SLC25 Mitochondrial Carrier Family: Structure and Mechanism. Trends Biochem Sci 45, 244–258 (2020).

37. Dzeja, P.P., Vitkevicius, K.T., Redfield, M.M., Burnett, J.C. & Terzic, A. Adenylate kinase-catalyzed phosphotransfer in the myocardium : increased contribution in heart failure. Circ Res 84, 1137–1143 (1999).

38. Pucar, D. et al. Cellular energetics in the preconditioned state: protective role for phosphotransfer reactions captured by 18O-assisted 31P NMR. J Biol Chem 276, 44812–44819 (2001).

39. Burkart, A., Shi, X., Chouinard, M. & Corvera, S. Adenylate kinase 2 links mitochondrial energy metabolism to the induction of the unfolded protein response. J Biol Chem 286, 4081–4089 (2011).

40. Six, E. et al. AK2 deficiency compromises the mitochondrial energy metabolism required for differentiation of human neutrophil and lymphoid lineages. Cell Death Dis 6, e1856 (2015).

41. Single, B., Leist, M. & Nicotera, P. Simultaneous release of adenylate kinase and cytochrome c in cell death. Cell Death Differ 5, 1001–1003 (1998).

42. Kohler, C. et al. Release of adenylate kinase 2 from the mitochondrial intermembrane space during apoptosis. FEBS Lett 447, 10–12 (1999).

43. Lee, H.J. et al. AK2 activates a novel apoptotic pathway through formation of a complex with FADD and caspase-10. Nat Cell Biol 9, 1303–1310 (2007).

44. Zhang, S. et al. Adenylate kinase AK2 isoform integral in embryo and adult heart homeostasis. Biochem Biophys Res Commun 546, 59–64 (2021).

45. Lagresle-Peyrou, C. et al. Human adenylate kinase 2 deficiency causes a profound hematopoietic defect associated with sensorineural deafness. Nat Genet 41, 106–111 (2009).

46. Pannicke, U. et al. Reticular dysgenesis (aleukocytosis) is caused by mutations in the gene encoding mitochondrial adenylate kinase 2. Nat Genet 41, 101–105 (2009).

47. Romero-Tamayo, S. et al. W196 and the beta-Hairpin Motif Modulate the Redox Switch of Conformation and the Biomolecular Interaction Network of the Apoptosis-Inducing Factor. Oxid Med Cell Longev 2021, 6673661 (2021).

48. Stocchi, V. et al. Simultaneous extraction and reverse-phase high-performance liquid chromatographic determination of adenine and pyridine nucleotides in human red blood cells. Anal Biochem 146, 118–124 (1985).

49. Lu, W., Wang, L., Chen, L., Hui, S. & Rabinowitz, J.D. Extraction and Quantitation of Nicotinamide Adenine Dinucleotide Redox Cofactors. Antioxid Redox Signal 28, 167–179 (2018).

50. Sallin, O. et al. Semisynthetic biosensors for mapping cellular concentrations of nicotinamide adenine dinucleotides. Elife 7 (2018).

51. Cambronne, X.A. et al. Biosensor reveals multiple sources for mitochondrial NAD(+). Science 352, 1474–1477 (2016).

52. Yang, H. et al. Nutrient-sensitive mitochondrial NAD+ levels dictate cell survival. Cell 130, 1095–1107 (2007).

53. Finger, Y. et al. Proteasomal degradation induced by DPP9-mediated processing competes with mitochondrial protein import. EMBO J 39, e103889 (2020).

54. Soboll, S., Scholz, R. & Heldt, H.W. Subcellular metabolite concentrations. Dependence of mitochondrial and cytosolic ATP systems on the metabolic state of perfused rat liver. Eur J Biochem 87, 377–390 (1978).

55. Ran, F.A. et al. Genome engineering using the CRISPR-Cas9 system. Nat Protoc 8, 2281–2308 (2013).

56. Stroud, D.A. et al. Accessory subunits are integral for assembly and function of human mitochondrial complex I. Nature 538, 123–126 (2016).

57. Montague, T.G., Cruz, J.M., Gagnon, J.A., Church, G.M. & Valen, E. CHOPCHOP: a CRISPR/Cas9 and TALEN web tool for genome editing. Nucleic Acids Res 42, W401–407 (2014).

58. Kuttner, V. et al. Global remodelling of cellular microenvironment due to loss of collagen VII. Mol Syst Biol 9, 657 (2013).

59. Cox, J. & Mann, M. MaxQuant enables high peptide identification rates, individualized p.p.b.-range mass accuracies and proteome-wide protein quantification. Nat Biotechnol 26, 1367–1372 (2008).

60. Wischhof, L. et al. Unbiased proteomic profiling reveals the IP3R modulator AHCYL1/IRBIT as a novel interactor of microtubule-associated protein tau. J Biol Chem 298, 101774 (2022).

61. Erdogan, A.J. et al. The mitochondrial oxidoreductase CHCHD4 is present in a semi-oxidized state in vivo. Redox Biol 17, 200–206 (2018).

62. Poepsel, S., Kasinath, V. & Nogales, E. Cryo-EM structures of PRC2 simultaneously engaged with two functionally distinct nucleosomes. Nat Struct Mol Biol 25, 154–162 (2018).

63. Punjani, A., Rubinstein, J.L., Fleet, D.J. & Brubaker, M.A. cryoSPARC: algorithms for rapid unsupervised cryo-EM structure determination. Nat Methods 14, 290–296 (2017).

64. Bepler, T. et al. Positive-unlabeled convolutional neural networks for particle picking in cryo-electron micrographs. Nat Methods 16, 1153–1160 (2019).

65. Emsley, P., Lohkamp, B., Scott, W.G. & Cowtan, K. Features and development of Coot. Acta Crystallogr D Biol Crystallogr 66, 486–501 (2010).

66. Liebschner, D. et al. Macromolecular structure determination using X-rays, neutrons and electrons: recent developments in Phenix. Acta Crystallogr D Struct Biol 75, 861–877 (2019).

67. Sanchez-Garcia, R. et al. DeepEMhancer: a deep learning solution for cryo-EM volume post-processing. Commun Biol 4, 874 (2021).

68. Pettersen, E.F. et al. UCSF ChimeraX: Structure visualization for researchers, educators, and developers. Protein Sci 30, 70–82 (2021).

69. Williams, C.J. et al. MolProbity: More and better reference data for improved all-atom structure validation. Protein Sci 27, 293–315 (2018).

70. Perez-Riverol, Y. et al. The PRIDE database resources in 2022: a hub for mass spectrometry-based proteomics evidences. Nucleic Acids Res 50, D543–D552 (2022).

