## Supplemental Information complete for "Interaction with AK2A links AIFM1 to cellular energy metabolism"

### TABLE OF CONTENT

#### SUPPLEMENTAL TABLES


Table S1. Cell lines page 3

Table S2. Primers and Plasmids page 5

Table S3. Antibodies page 6

Table S4. Further Tools and Equipment page 6

Table S5. Software and Algorithms page 6


#### SUPPLEMENTAL FIGURES


Figure S1 page 7

Figure S2 page 9

Figure S3 page 10

Figure S4 page 11

Figure S5 page 12

Figure S6 page 13

Figure S7 page 14

Figure S8 page 15

Figure S9 page 16

Figure S10 page 17

Figure S11 page 18

Figure S12 page 20

Figure S13 page 22

Figure S14 page 24

Figure S15 page 25

### SUPPLEMENTAL TABLES

#### Table S1. Cell lines

| Cell line | Plasmid | Gene | Tag | Reference |
|---|---|---|---|---|
| **Flp-In T-Rex-293** | -- | -- | -- | ThermoFisher, R78007 |
| **Flp-In T-Rex-293 AIFM1 knockout #5-AIFM1** | pcDNA5/FRT/TO | ORF AIFM1 | HA (C-terminal) | This work |
| **Flp-In T-Rex-293 AIFM1 knockout #5-mock** | pcDNA5/FRT/TO | -- | -- | This work |
| **Flp-In T-REx-293 AK2 knockout #2.14-AK2A** | pcDNA5/FRT/TO | ORF AK2A | -- | This work |
| **Flp-In T-REx-293 AK2 knockout #2.14-AK2B** | pcDNA5/FRT/TO | ORF AK2B | -- | This work |
| **Flp-In T-REx-293 AK2 knockout #2.14-mock** | pcDNA5/FRT/TO | -- | -- | This work |
| **Flp-In T-REx-293-MIA40** | pcDNA5/FRT/TO | ORF *MIA40* | HA (C-terminal) | [1] |
| **Flp-In T-REx-293-Mock** | pcDNA5/FRT/TO | -- | -- | [1] |
| **HEK293T** | -- | -- | -- | [2] |
| **HEK293T AIFM1 knockout #9** | -- | Deletion of AIFM1 | -- | [3] |
| **HEK293T AIFM1 knockout #9-AIFM1** | PB-CuO-MCSIRES-GFP-EF1-CymR-Puro | ORF AIFM1 | HA (C-terminal) | [3] |
| **HEK293T AIFM1 knockout #9-SMACmtsAIFM1** | PB-CuO-MCSIRES-GFP-EF1-CymR-Puro | ORF SMAC (1-???) ORF AIFM1 (???-613) | HA (C-terminal) | This work |
| **HEK293T AIFM1 knockout #9-mock** | PB-CuO-MCSIRES-GFP-EF1-CymR-Puro | -- | -- | [3] |
| **HEK293T-MIA40** | PB-CuO-MCSIRES-GFP-EF1-CymR-Puro | ORF MIA40 | -- | [1] |
| **HEK293T-Mock** | PB-CuO-MCSIRES-GFP-EF1-CymR-Puro | -- | -- | [3] |
| **HeLa** | -- | -- | -- | -- |
| **HepG2** | -- | -- | -- | -- |
| **C2C12** | -- | -- | -- | Sigma (cat. 91031101) |
| **Rosetta2 (DE3)-AIFM1** | pET-24a(+) | ORF AIFM1 *(103-613)* | 6xHis (C-terminal) | [3] |

| | | | | |
|---|---|---|---|---|
| **Rosetta2 (DE3)-AK2A** | pET-15(b) | ORF AK2A *($C_{40}$▶S; $C_{232}$▶S)* | 6xHis (N-terminal) | This work |
| **Rosetta2 (DE3)-AK2B** | pET-15(b) | ORF AK2B *($C_{40}$▶S)* | 6xHis (N-terminal) | This work |
| **Rosetta2 (DE3)-MIA40 SPS** | pET-24a(+) | ORF MIA40 *($C_{4}$▶S; $C_{53}$▶S; $C_{55}$▶S)* | 6xHis (C-terminal) | This work |

### Table S2. Primers and Plasmids

| Plasmid | Primer (5´-3´) | Restriction sites | Reference |
|---|---|---|---|
| **AIFM1 in PB-CuO-MCSIRES-GFP-EF1-CymR-Puro** | FW: GCTTCGAAGGCATGTTCCGGTGTGGAG<br>RV: CCGCGGCCGCTCAAGCGTAATCTGGAACATCGTATGGGTAGTCTTCATGAATGTTG | Bsp119<br>NotI | [3] |
| **AIFM1 in pcDNA5/FRT/TO** | FW: GCCGGTACCGAAATGTTCCGGTGTGGAGGC<br>RV: GTTGCGGCCGCTCAAGCATAATCTGGAACATCATATGGATAACTTCCGTCTTCATGAATGTTGAATAGTTTGGC | KpnI<br>NotI | [3] |
| **AIFM1 in pET-24(a)** | FW: CGCGAATTCATGGGGCTGACACCAG<br>RV: GCGGCGGCCGCGTCTTCATGAATGTTG | EcoRI<br>NotI | [3] |
| **AK2A in pcDNA5/FRT/TO** | FW: GCGCATATGGCTCCCAGC<br>RV: CGCGCTCAGCTTAGATAAACATAACCAAGTCTTTAC | NdeI<br>Bsu1102I | This work |
| **AK2A in pET-15(b)** | FW: GCGCATATGGCTCCCAGC<br>RV: CGCGCGGCCGCTTAGATAAACATAACCAAGTCTTTAC | NdeI NotI | This work |
| **AK2B in pcDNA5/FRT/TO** | FW: GCGCATATGGCTCCCAGC<br>RV: CGCGCTCAGCCTAGGATGTGGCTTT | NdeI<br>Bsu1102I | This work |
| **AK2B in pET-15(b)** | FW: GCGCATATGGCTCCCAGC<br>RV: CGCGCGGCCGCTTACTAGGATGTGGCTTTG | NdeI NotI | This work |
| **MIA40 in PB-CuO-MCSIRES-GFP-EF1-CymR-Puro** | FW: GCCGCTAGCGCAGCCATGTCCTATTGCCGGCAGGAAG<br>RV: GTAGCGGCCGCTTATTTCTCAAATTGTGGATGACTCCATCCTCCAGCACTTGATCCCTCCTCTTCTTTG | NheI<br>NotI | [1] |
| **MIA40 SPS in pET-24a(+)** | FW: CGCCATATGTCCTATTGCC<br>RV: CGCCTCGAGCTCATCCTCTTG | NdeI<br>XhoI | This work |
| **PB-CuO-MCSIRES-GFP-EF1-CymR-Puro empty vector** | System Biosciences Cat#PBQM812A-1 | -- | -- |
| **pcDNA5/FRT/TO empty vector** | Invitrogen V652020 | -- | -- |
| **Myc-DDK-AK2** | Origene, cat. RC210614 | | |
| **Myc-DDK-MIC27 Plasmid** | Origene, cat. RC214060 | | |
| **Super PiggyBac Transposase Expression vector** | System Biosciences Cat#PB210PA-1 | -- | -- |

### Table S3. Antibodies

| Antibody | Company | Identifier |
|---|---|---|
| **Goat anti-Mouse IgG (H&L), HRP Conjugate** | ImmunoReagents | Cat# GtxMu-003-DHRPX |
| **Goat anti-Rabbit IgG (H&L), HRP Conjugate** | ImmunoReagents | Cat# GtxRb-003-DHRPX |
| **Rabbit polyclonal anti-AIFM1** | Chemicon | Cat# ab16501 |
| **Rabbit polyclonal anti-AK2** | (Finger *et al.*, 2020) | N/A |
| **Rabbit polyclonal anti-MIC19** | Proteintech | Cat# 25625-1-AP |
| **Rabbit polyclonal anti-MIC27** | Proteintech | Cat# 28514-1-AP |
| **Rabbit polyclonal anti-CPOX** | St John's Laboratory | Cat# STJ23214 |
| **Rabbit polyclonal anti-HA** | Sigma-Aldrich | Cat# SAB4300603 |
| **Rabbit polyclonal anti-MIA40** | [4] | N/A |

### Table S4. Further Tools and Equipment

| | Company/ Source | Identifier |
|---|---|---|
| **cycloheximide** | Sigma | Cat# 239763 |
| **FuGENE HD Transfection Reagent** | Promega | Cat# E2311 |
| **Methyl-PEG-Maleimide, mmPEG24** | Thermo Fisher | Cat# 22713 |
| **One Shot TOP10 Chemically Competent *E. coli*** | Thermo Fisher | Cat# C404010 |
| **Pierce 660 nm Protein Assay Reagent** | Thermo Scientific | Cat# 22660 |
| **Rosetta™ 2 (DE3) Singles™ Competent Cells** | Novagen | Cat# 70954-3 |
| **ROTI®Quant universal** | Carl Roth | Cat# 0120.1 |
| **UltrAuFoil® R 1.2/1.3 grid** | Quantifoil | Cat# N1-A14nAu30-01 |

### Table S5. Software and Algorithms

| | Company/ Source | Identifier |
|---|---|---|
| **ChimeraX (version 1.7.1)** | | https://www.cgl.ucsf.edu/chimerax |
| **Coot (version 0.9.8.7)** | [5] | |
| **Corel draw** | Corel Corporation | |
| **CryoSparc (version 4.4)** | [6] | |
| **Image Lab 5.2** | Biorad Laboratories | |
| **Phenix (version 1.21)** | [7] | |

### SUPPLEMENTARY FIGURES

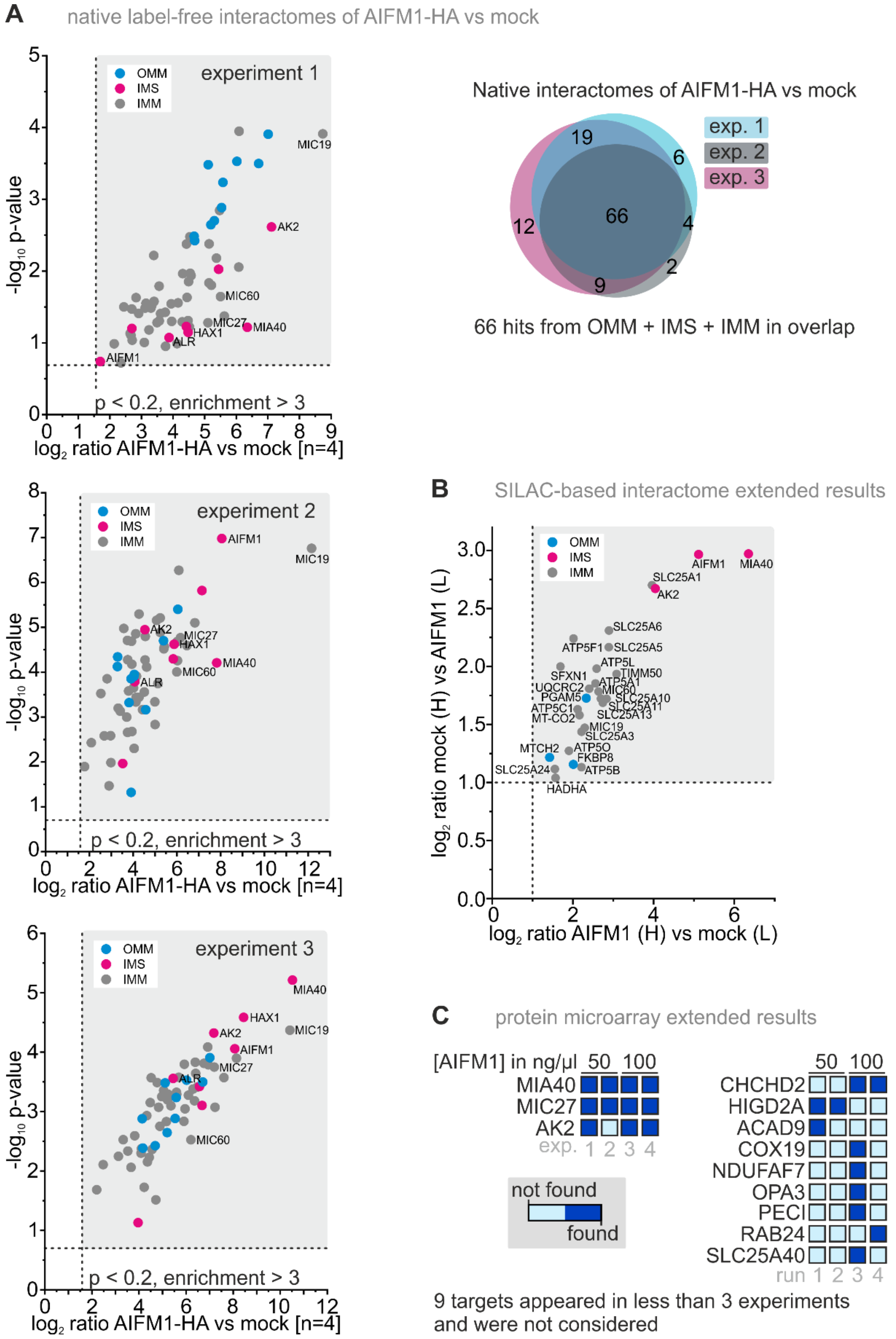


**Supplementary Figure S1 (linked to Figure 1). A high-confidence interactome reveals AK2 and MICOS components as novel AIFM1 interaction partners.**

**(A)** Individual data sets for the three repeats with four biological replicates each shown in **Figure 1C**. The interactomes show considerable overlap leading to the identification of 66 potential interactors of AIFM1-HA.

**(B)** SILAC-based data set for the experiment shown in **Figure 1D**. We identified 27 potential interactors localized in IMM, IMS and OMM. Notably, the datasets from (A) and (B) do not only show AK2, MIA40, and MICOS components as potential AIFM1 interactors but also members of the SLC25 family (including the ADP/ATP carrier SLC25A5) and the ATPase (including ATP5A1, ATP5B, ATP5F1, ATP5L, and ATP5O)

**(C)** Extended results for the protein microarray shown in **Figure 1E**. Many different targets were only identified in single experiments. Only AK2, MIA40 and MIC27 were consistently identified.

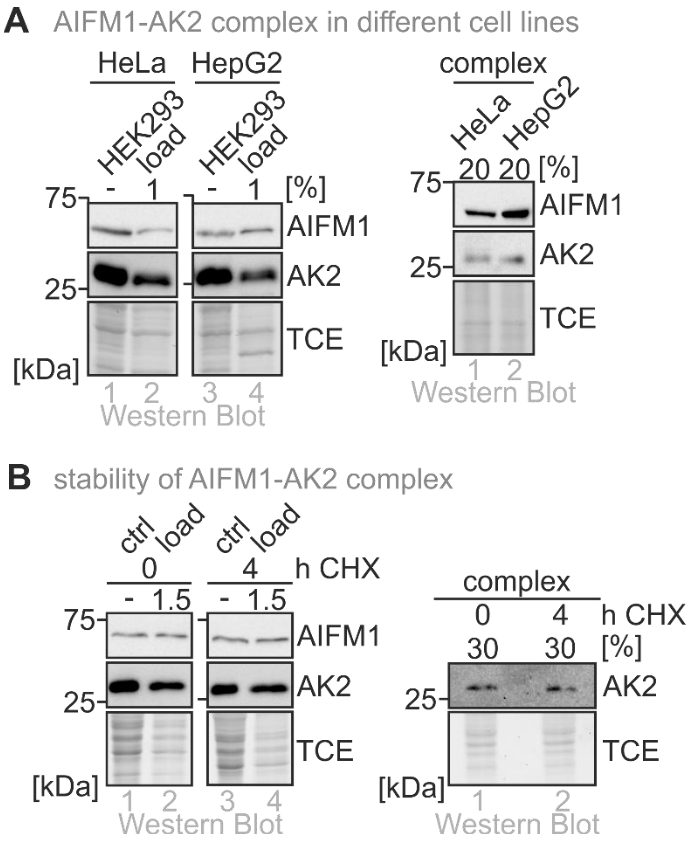


**Supplementary Figure S2 (linked to Figure 2). Properties of the AIFM1-AK2 complex.**

**(A)** The AIFM1-AK2A complex is present in different cell lines. Experiment performed as in **Figure 2F** in the indicated cell lines.

**(B)** The AIFM1-AK2A complex is stable. Experiment performed as in **Figure 2F** except that cells were treated with the translation inhibitor cycloheximide for 4 hours or left untreated. The amounts of AK2A in the AIFM1-AK2A complex do not change during this time indicating a stable complex.

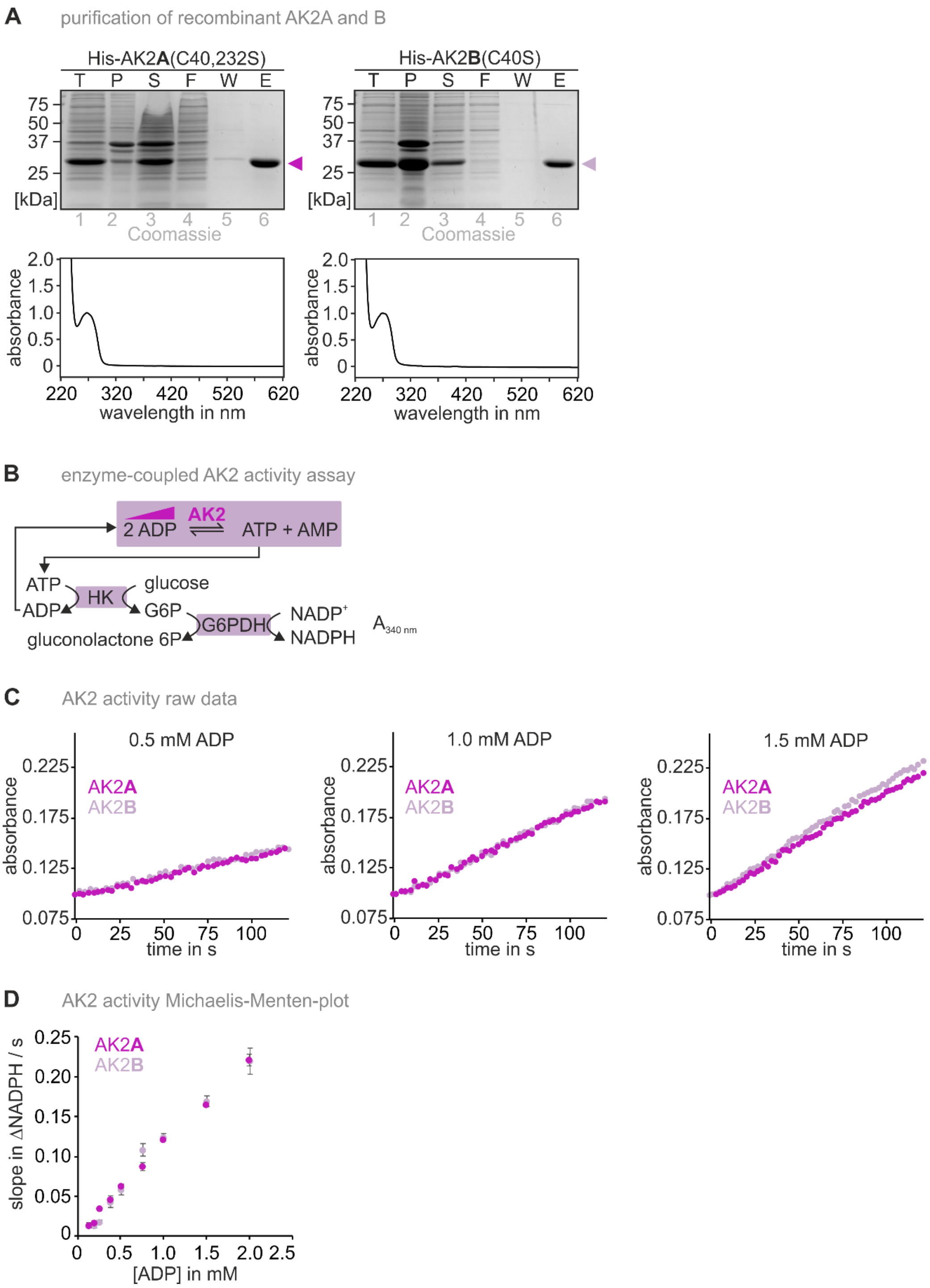


**Supplementary Figure S3 (linked to Figure 2). Recombinant AK2 isoform A and B exhibit a similar enzymatic profile.**

**(A)** Purification of the isoforms His-AK2A and His-AK2B. Both proteins were well-behaved and can be purified in similar amounts and to similar purity. T: total, P: pellet, S: supernatant, F: flow through, W: wash, E: eluate

**(B)** AK2 activity assay with varying ADP concentrations. Hexokinase (HK) and glucose-6-phosphate dehydrogenase (G6PDH) couple the reduction of $NADP^+$ to the interconversion of adenine nucleotides. G6P, glucose-6-phosphate

**(C,D)** Raw data **(C)** and velocity *vs* ADP concentration slope plot **(D)** for the enzyme activities of AK2A and AK2B. The concentration of ADP is titrated. Both isoforms exhibit the same activity towards ADP.

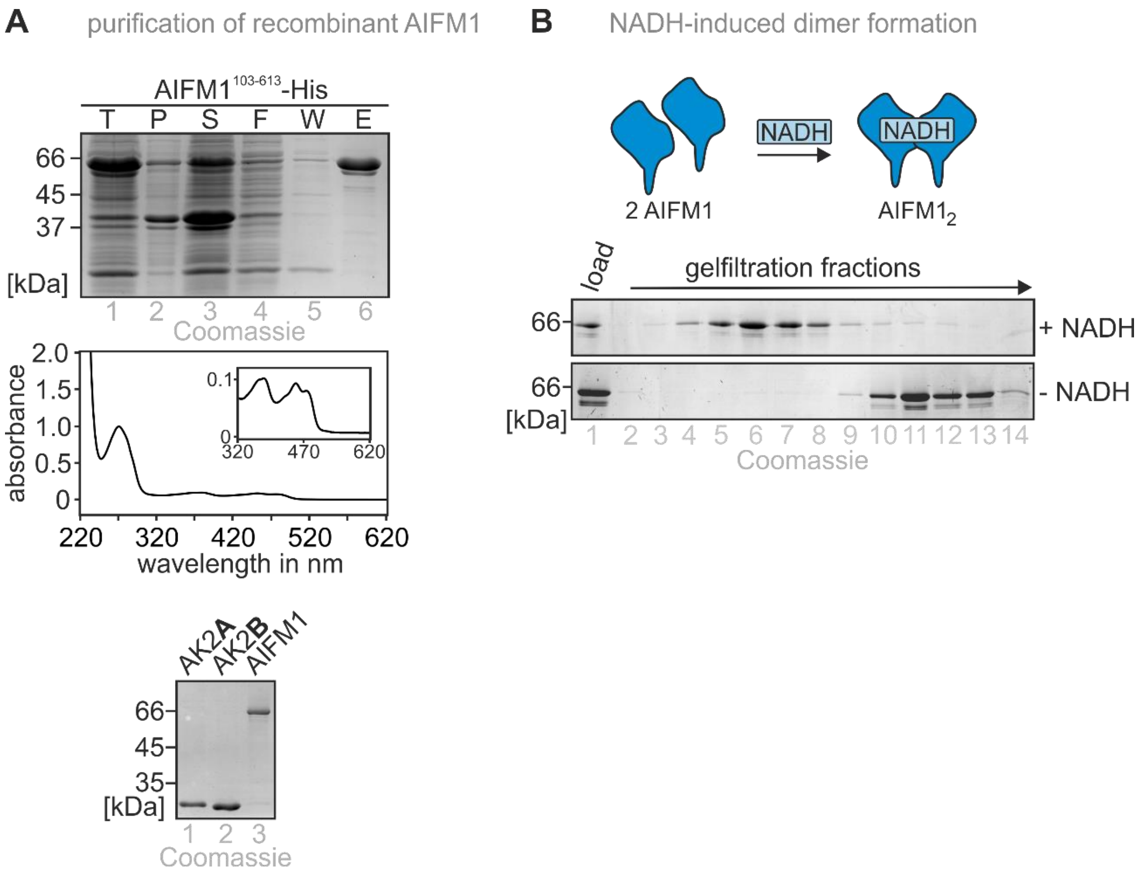


**Supplementary Figure S4 (linked to Figure 2). NADH-induced *in vitro* dimerization of recombinant soluble AIFM1.**

**(A)** Purification of soluble AIFM1 (AIFM1 103-613). AIFM1 lacking the mitochondrial targeting signal and the transmembrane domain was purified and contains the FAD cofactor. Purified AK2A and AK2B used in the *in vitro* reconstitution assay of the AIFM1-AK2 complex were loaded for comparison onto the same gel as AIFM1. T: total, P: pellet, S: supernatant, F: flow through, W: wash, E: eluate

**(B)** NADH-induced dimer formation of AIFM1. NADH addition leads to rapid AIFM1 dimerisation that can be followed by gel filtration. In the absence of NADH, AIFM1 migrates at the height of the monomer.

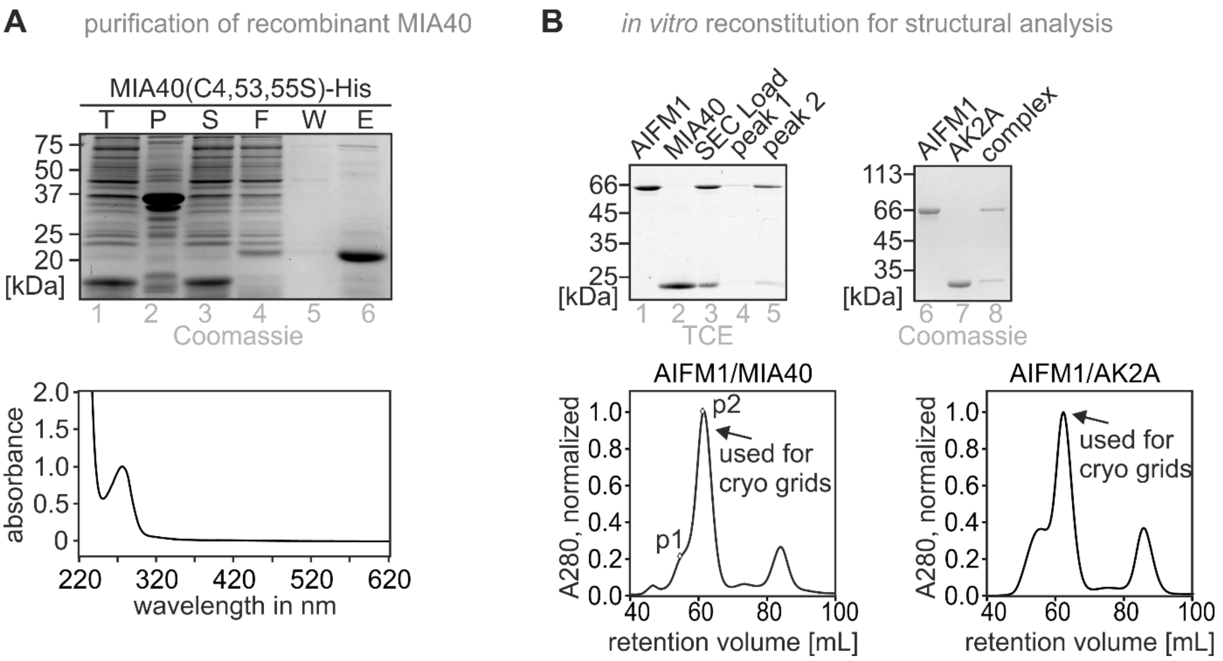


**Supplementary Figure S5 (linked to Figure 3). Protein preparation for cryo-EM.**

**(A)** Purification of the redox-inactive MIA40-C4,53,55S variant. T: total, P: pellet, S: supernatant, F: flow through, W: wash, E: eluate

**(B)** *In vitro* reconstitution of the AIFM1-MIA40 and AIFM1-AK2A complexes and isolation of the complexes by gel filtration for cryo-EM.

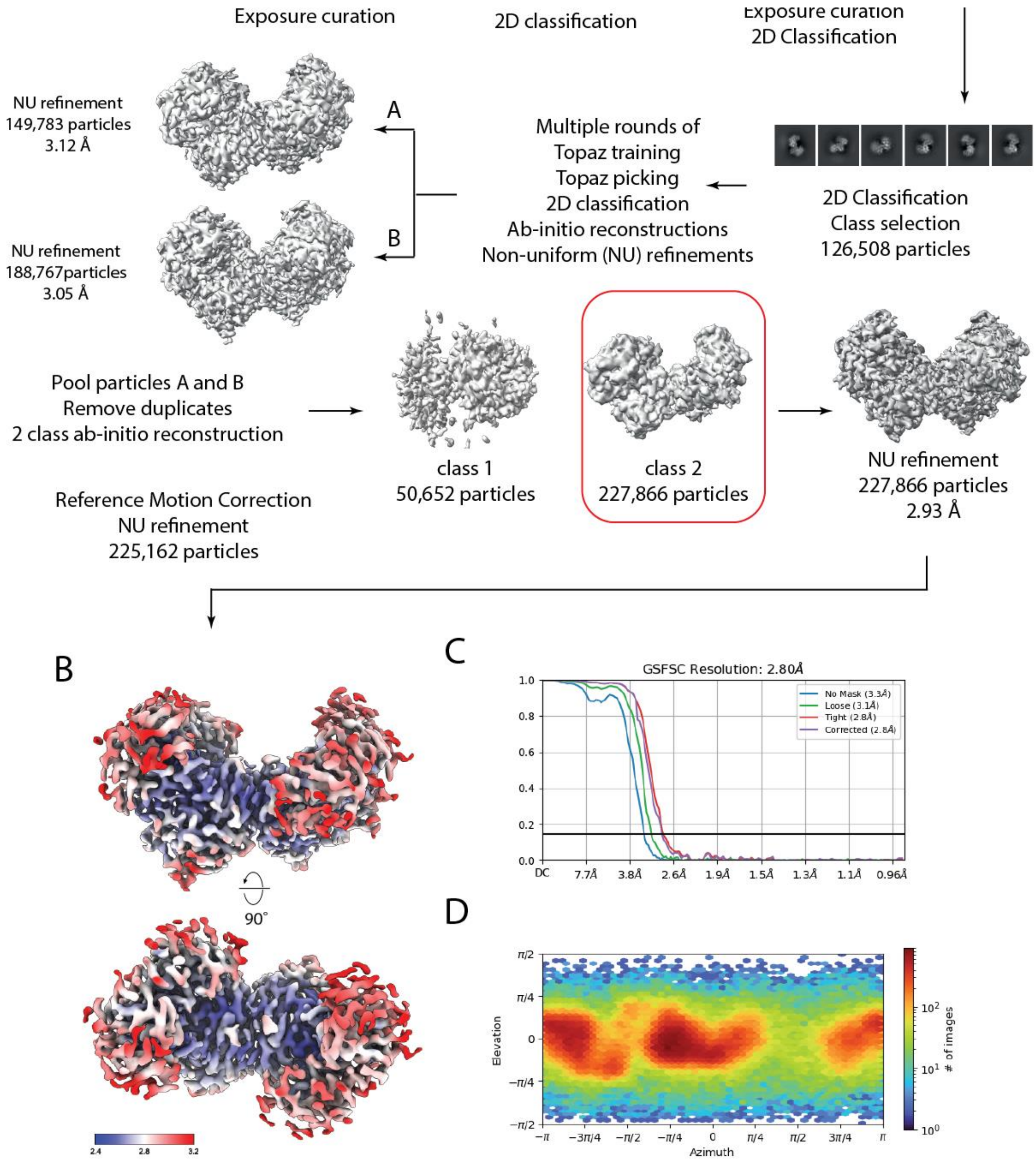


**Supplementary Figure S6 (linked to Figure 3). Cryo-EM data processing workflow of the AIFM1 dimer.**

**(A)** Processing workflow for the AIFM1 dimer. All data processing was performed in cryoSPARC [6].

**(B)** Final reconstruction colored according to local resolution, ranging from 2.4 Å (blue) to 3.2 Å (red).

**(C)** The global resolution determined by the FSC cut-off at 0.143 was 2.8 Å.

**(D)** Euler angle distribution plot.

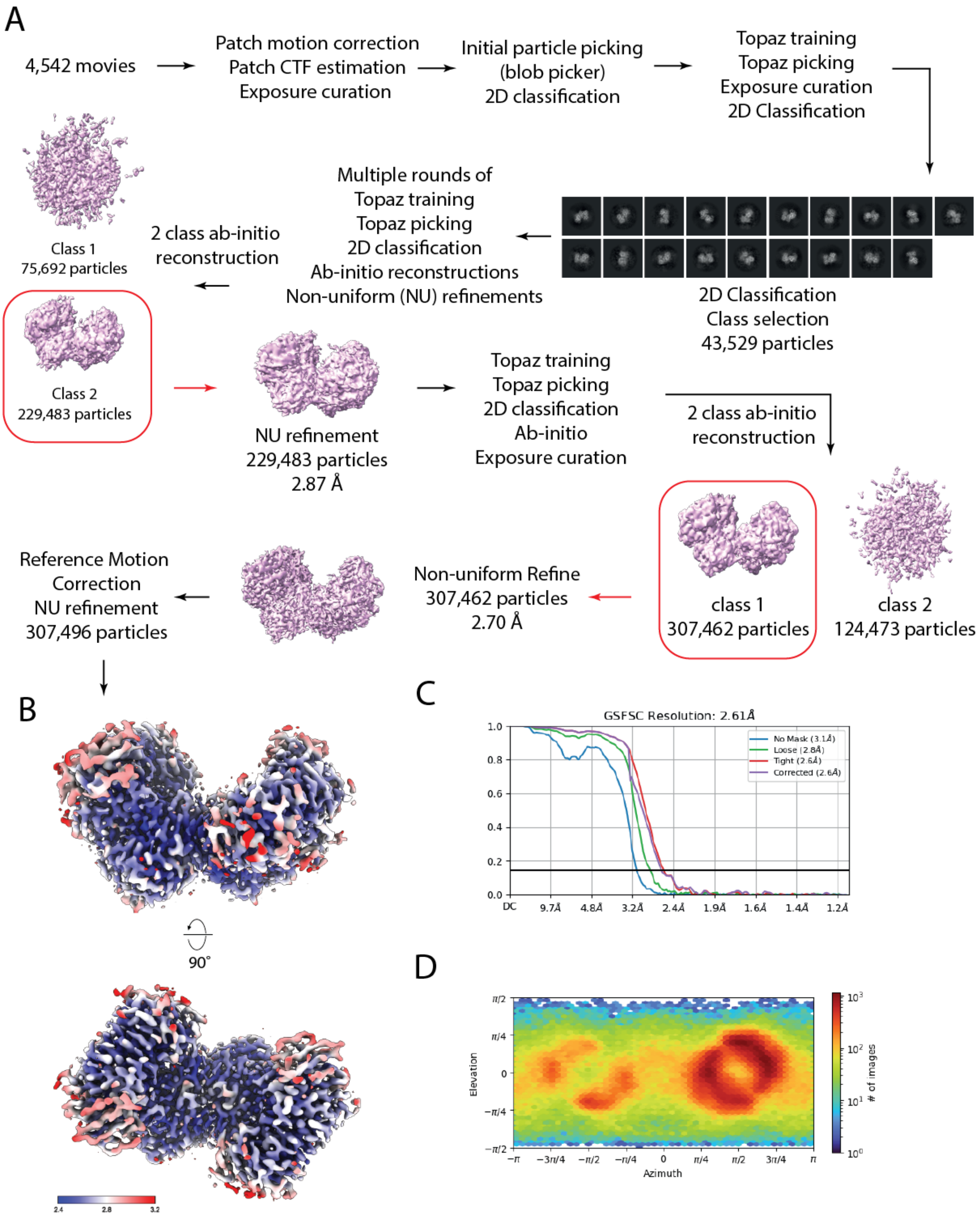


**Supplementary Figure S7 (linked to Figure 3). Cryo-EM data processing workflow of the AIFM1-AK2A complex.**

**(A)** Processing workflow for the AIFM1-AK2A complex. All data processing was performed in cryoSPARC [6].

**(B)** Final reconstruction colored according to local resolution, ranging from 2.4 Å (blue) to 3.2 Å (red).

**(C)** The global resolution determined by the FSC cut-off at 0.143 was 2.6 Å.

**(D)** Euler angle distribution plot.

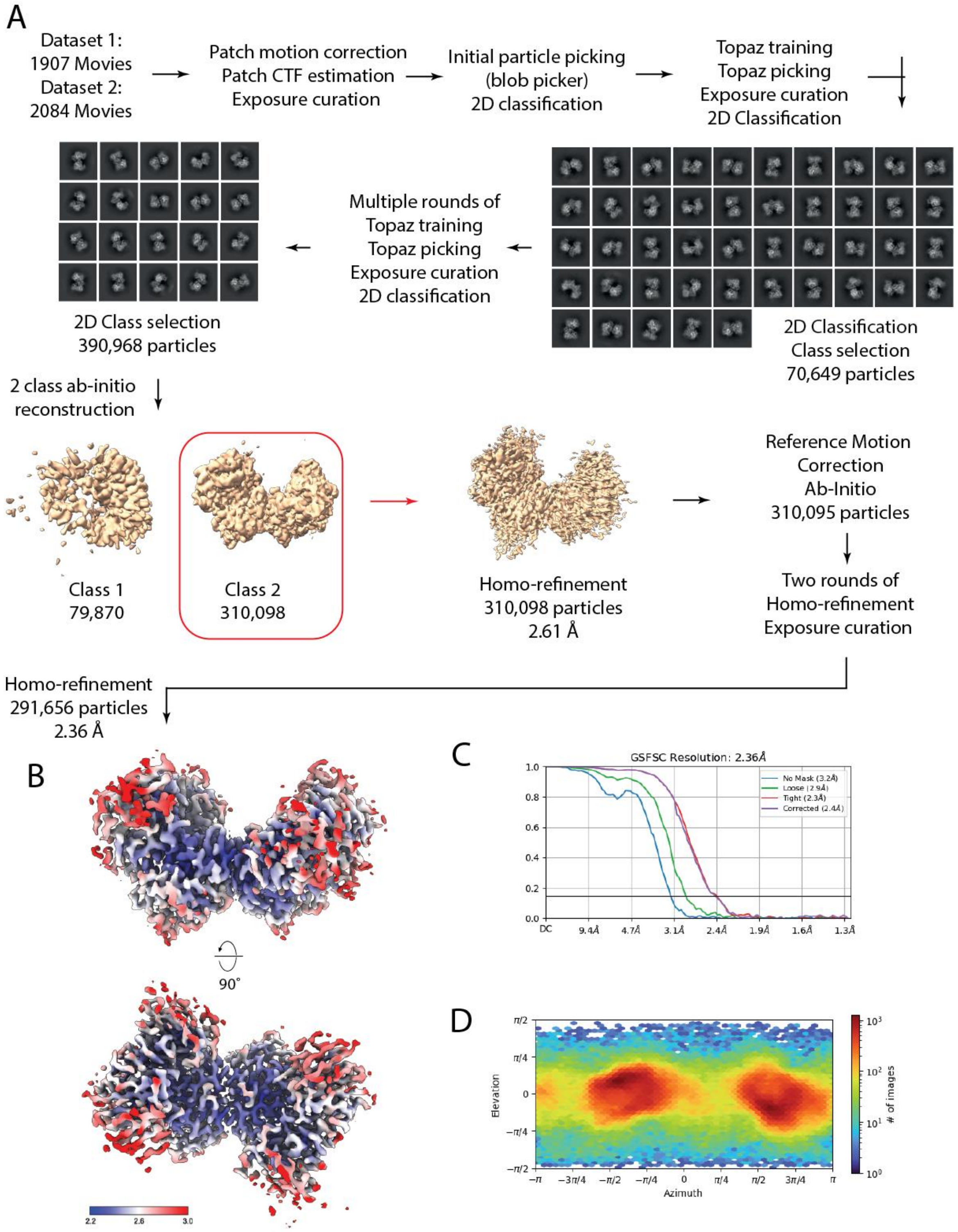


**Supplementary Figure S8 (linked to Figure 3). Cryo-EM data processing workflow of the AIFM1-MIA40 complex.**

**(A)** Processing workflow for the AIFM1-MIA40 complex. All data processing was performed in cryoSPARC [6].

**(B)** Final reconstruction colored according to local resolution, ranging from 2.2 Å (blue) to 3.0 Å (red).

**(C)** The global resolution determined by the FSC cut-off at 0.143 was 2.4 Å.

**(D)** Euler angle distribution plot.

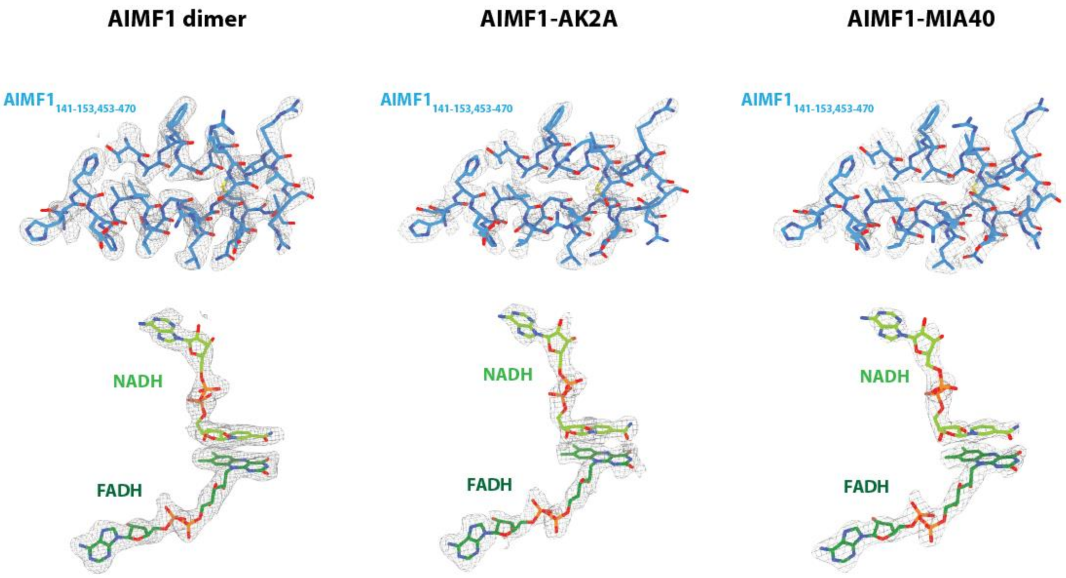


**Supplementary Figure S9 (linked to Figure 3). Cryo-EM map quality.**

Exemplary cryo-EM densities (grey mesh) and atomic models obtained in this study for AIFM1 (aa 141-153,453-470), top panels, or the NAD and FAD cofactors (bottom panels), of all three structures shown here.

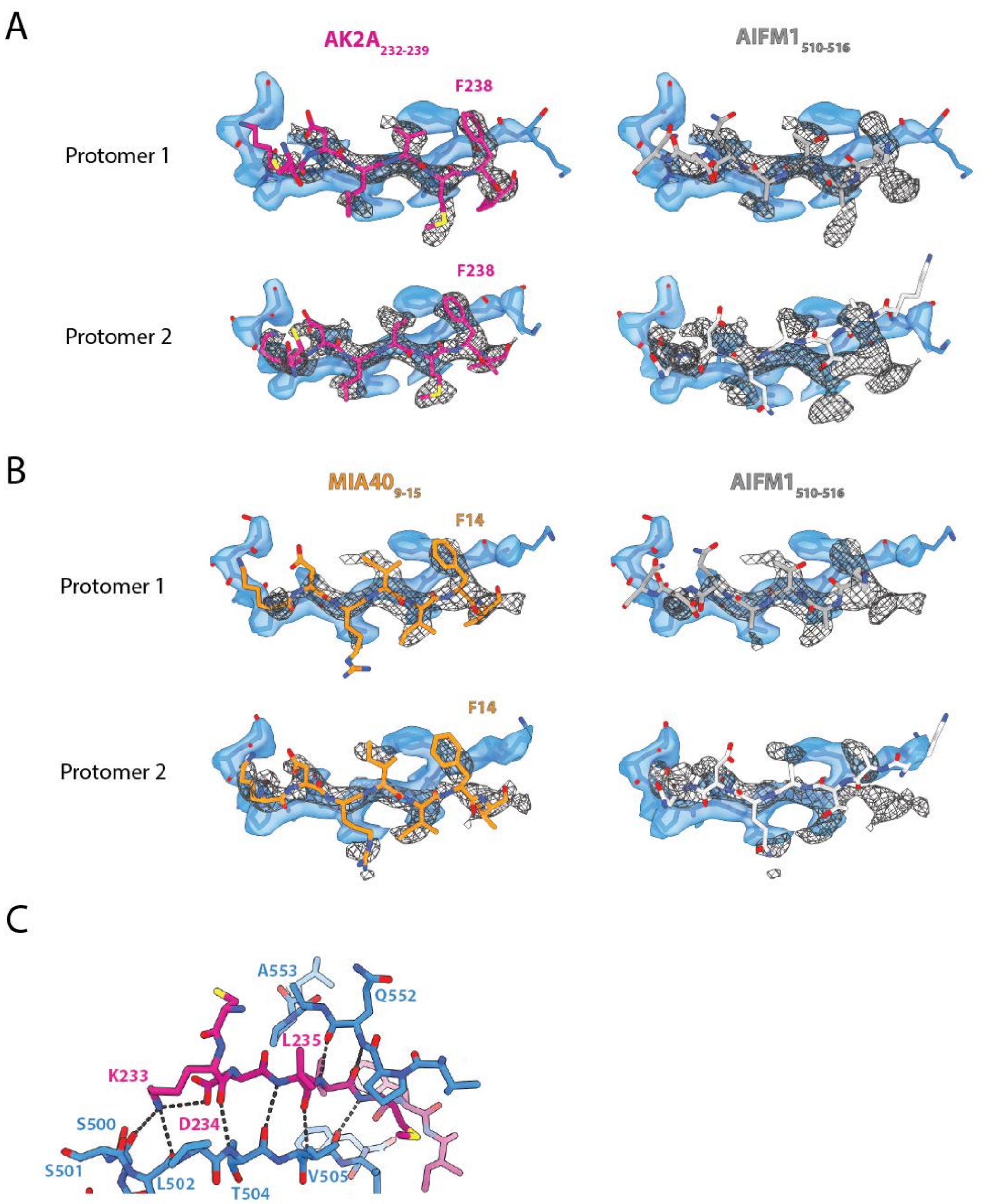


**Supplementary Figure S10 (linked to Figure 3). Details of AK2A and MIA40 interactions with AIFM1, unambiguous fit of AK2A and MIA40.**

**(A)** Cryo-EM map of the AIFM1-AK2A complex and fitted model for AIFM1 (aa 500-510), (blue, transparent map) and AK2A (aa 232-239), (purple, grey mesh map, left panels) showing the fit with AK2A as opposed to AIFM1 (aa 510-514), (grey model, right panels).

**(B)** Cryo-EM map of the AIFM1-MIA40 complex and fitted model for AIFM1 (aa 500-510), (blue, transparent map) and MIA40 (aa 9-15), (orange, grey mesh map, left panels) showing the fit with MIA40 as opposed to AIFM1 (aa 539-544), (grey model, right panels).

**(C)** Details of the interactions of AK2A (purple) with the C-loop of AIFM1 (blue), and hydrogen bonds between the conserved K233 and D234 of AK2A (corresponding to K9 and D10 of MIA40) and AIFM1.

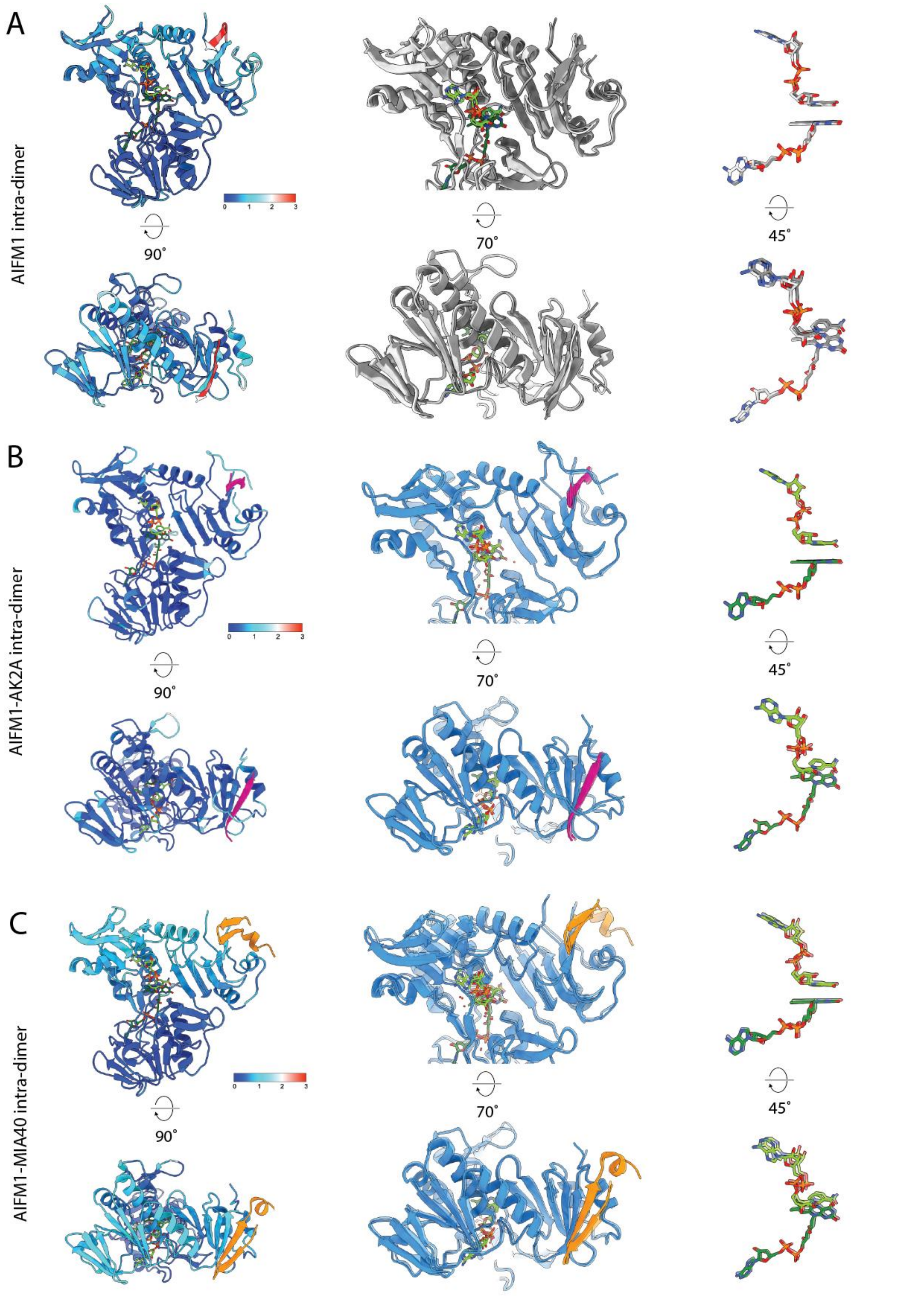


**Supplementary Figure S11 (linked to Figure 4). Structural variability between protomers within each AIFM1 dimer of each complex.**

AIFM1 monomers within each complex were superimposed using UCSF ChimeraX at N-terminal residues including the dimer interface (aa 232-257, 404-434 and 440-450). Left panels: models colored according to the root-mean-square deviation (rmsd) of the Cα atoms. Color code: blue = 0 Å, cyan = 1 Å, white = 2 Å, red = 3 Å. Middle panels: enlarged view of an overlay of both monomer models after alignment, one of the models transparent, to highlight displacement. Right panels: enlarged view of the NAD and FAD cofactors of both monomers after alignment of the models to visualize the direction and extent of variability. **(A)** AIFM1 dimer. **(B)** AIFM1-AK2A complex. **(C)** AIFM1-MIA40 complex.

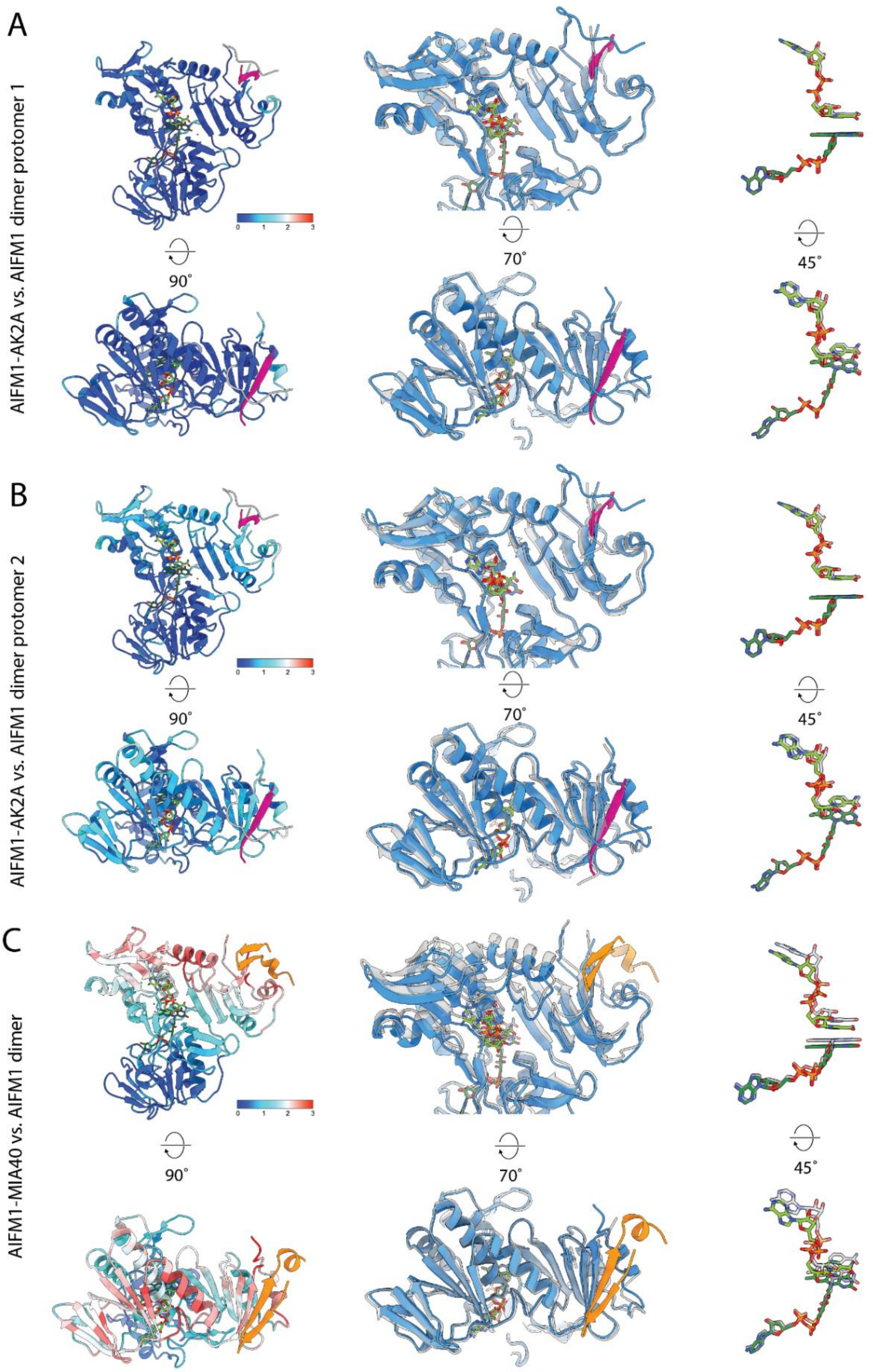


**Supplementary Figure S12 (linked to Figure 4). Structural variability between monomers of AIFM1-MIA40 and AIFM1-AK2A complexes as compared to the AIFM dimer.**

Monomers of the AIFM1-AK2A complex (**A,B**) or the AIFM1-MIA40 complex (**C**) were superimposed using UCSF ChimeraX at the N-terminal β-sheets (aa 128-165 and aa 212-261). Left panels: models coloured according to the root-mean-square deviation (rmsd) of the Cα atoms. Color code: blue = 0 Å, cyan = 1 Å, white = 2 Å, red = 3 Å. Middle panels: enlarged view of an overlay of both monomer models after alignment, one of the models transparent, to highlight displacement. Right panels: enlarged view of the NAD and FAD cofactors of both monomers after alignment of the models to visualize the direction and extent of variability. For AIFM1-AK2A, the alignment of one monomer to each of the AIFM1 dimer protomers is shown. For AIFM1-MIA40, only the monomer to monomer comparison with the strongest variability is shown.

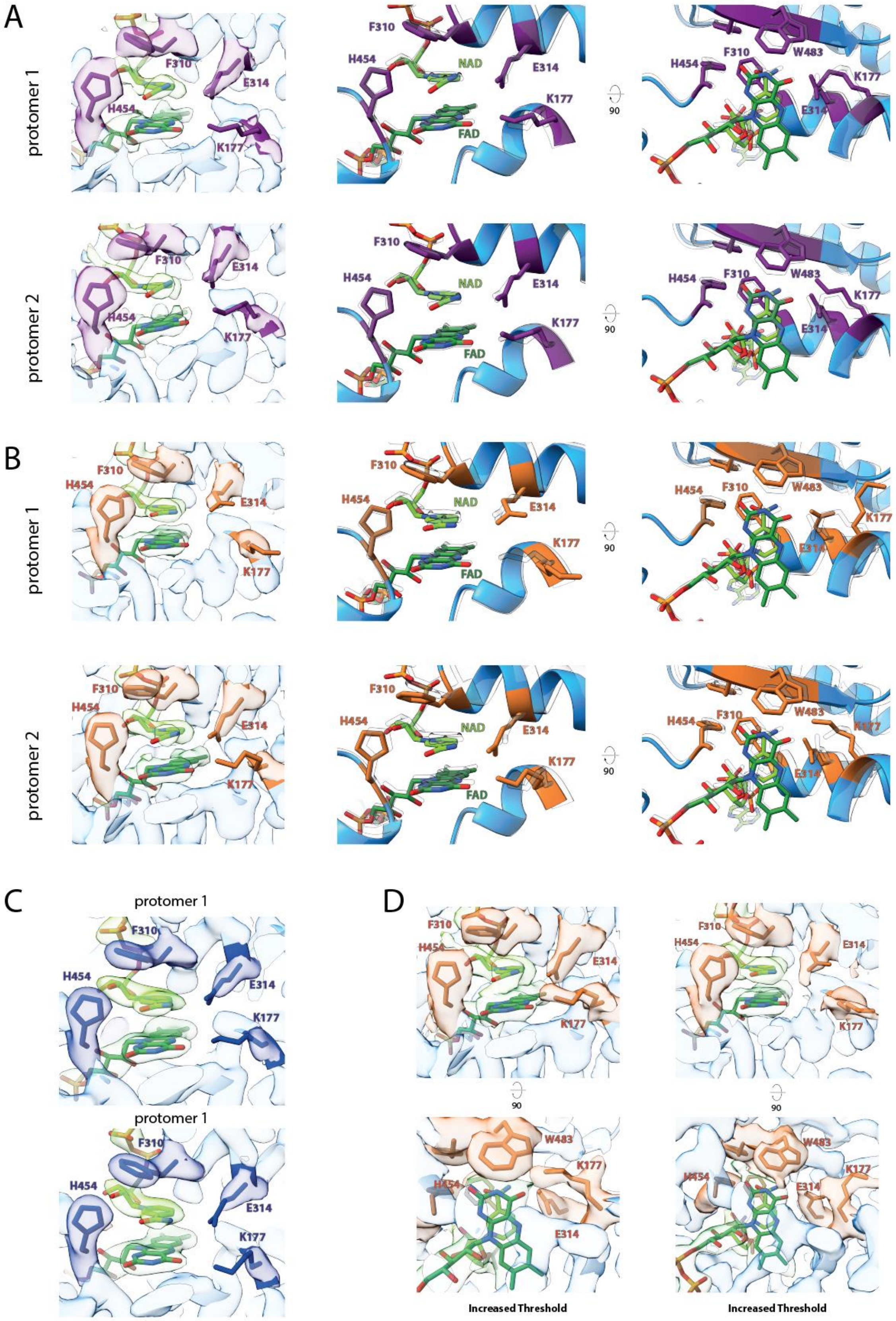


**Supplementary Figure S13 (linked to Figure 4). Structural details and conformational changes of the active site of AIFM1.**

**(A)** Active site of the two protomers (top and bottom row) of AIFM1-AK2A. Left panel: Local cryo-EM density (semi-transparent) and fitted model, cofactor binding residues shown as stick representation. Middle and right panels: Enlarged views of the AIFM1 active site, residues stabilizing the cofactors shown as stick representation and in purple. Model of the AIFM1 dimer shown transparent and as an overlay.

**(B)** Active site of the two protomers (top and bottom row) of MIA40-bound AIFM1. Left panel: Local cryo-EM density (semi-transparent) and fitted model, cofactor binding residues shown as stick representation. Middle and right panels: Enlarged views of the AIFM1 active site, residues stabilizing the cofactors shown as stick representation and in orange. Model of the AIFM1 dimer shown transparent and as an overlay.

**(C)** Cryo-EM density of the active site of the AIFM1 dimer of both protomers (top and bottom panels) with the respected atomic model fitted.

**(D)** Cryo-EM density of the active site of AIFM1-MIA40 of both protomers (left and right panels) shown at increased threshold (bottom panels) to visualize weaker densities indicating flexibility.

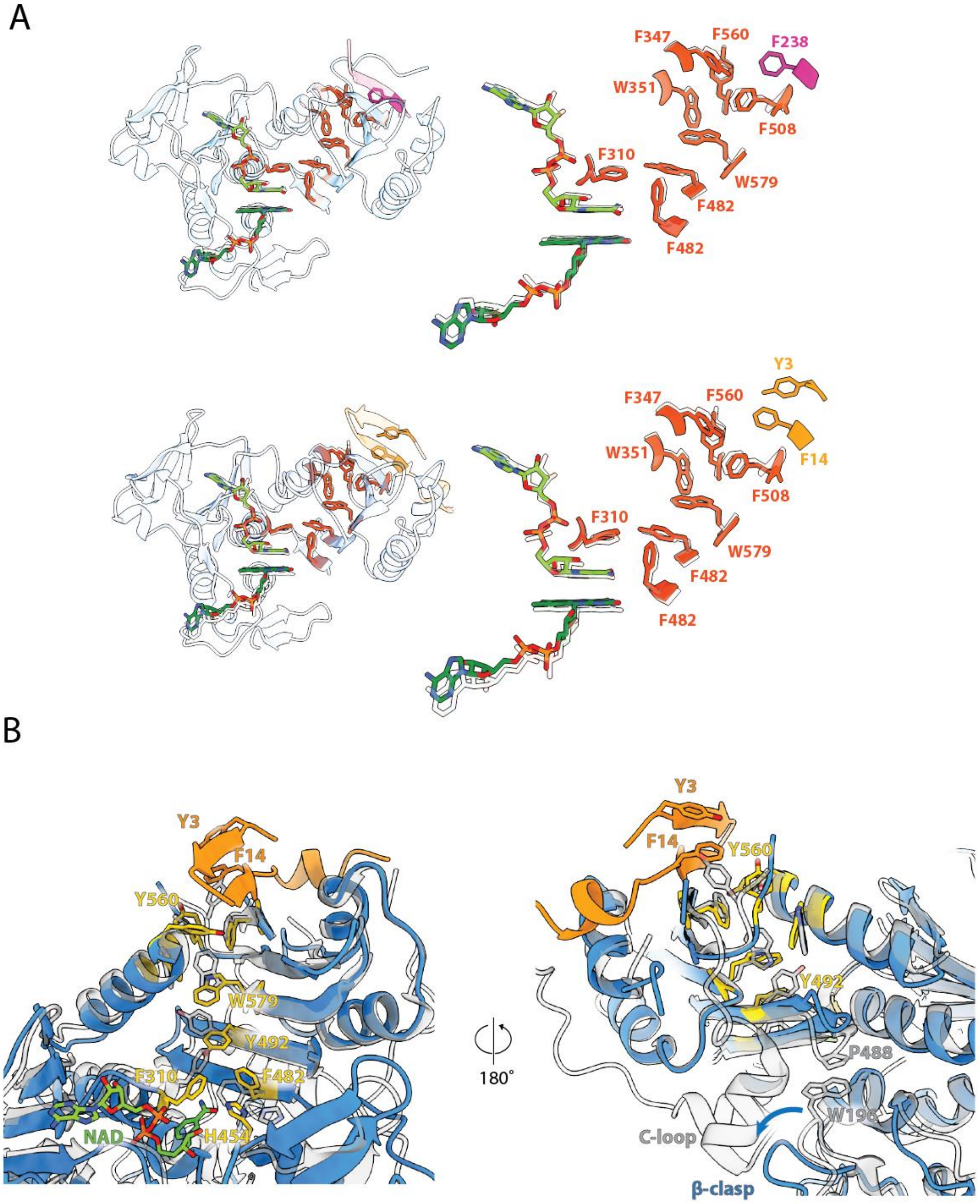


**Supplementary Figure S14 (linked to Figure 4). Aromatic tunnel of AIFM1 and conformational stabilization of the aromatic tunnel by AK2A and MIA40 binding.**

**(A)** Aromatic tunnel of AIFM1. Left panel: aromatic residues of AIFM1 (dark orange), AK2A (top, purple) and MIA40 (bottom, light orange) forming the aromatic tunnel within an AIFM1 monomer, linking the cofactor binding site and protein surface. Right panel: enlarged detail of amino acids and the NAD and FAD cofactors involved. Transparent model: overlay of the AIFM1 dimer lacking AK2A or MIA40 binding.

**(B)** Structural details of aromatic aa side chains forming the 'aromatic tunnel' and the conformational impact of MIA40 binding (orange). Aromatic tunnel residues and the NAD binding H454 are highlighted in yellow. The AIFM1 model in the monomeric, oxidized conformation (PDB 4BV6, [8]) is shown as a grey, transparent overlay.

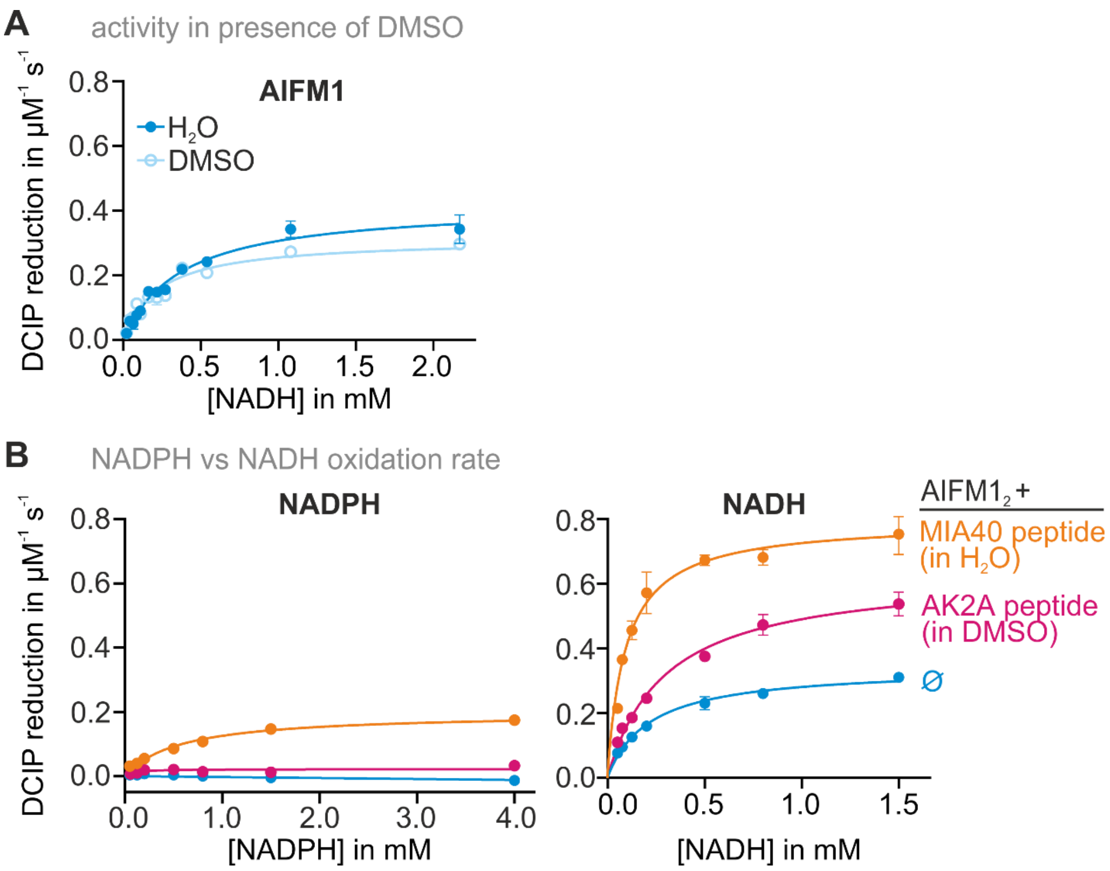


**Supplementary Figure S15 (linked to Figure 4). AIFM1 changes its activity upon binding of AK2 or MIA40.**

**(A)** Activity of AIFM1 towards NADH and DCIP does not change in the presence of low amounts of DMSO.

**(B)** Addition of AK2 or MIA40 to AIFM1 increase its activity towards NADH and DCIP. AIFM1 activity towards NADPH is in this assay negligible. Only binding of MIA40 to AIFM1 results in a minor increase of AIFM1 activity. Please observe the differences in the X-axis of the two plots in this panel.

### REFERENCES


1. Murschall, L.M. et al. The C-terminal region of the oxidoreductase MIA40 stabilizes its cytosolic precursor during mitochondrial import. *BMC Biol* **18**, 96 (2020).
2. Habich, M. et al. Vectorial Import via a Metastable Disulfide-Linked Complex Allows for a Quality Control Step and Import by the Mitochondrial Disulfide Relay. *Cell Rep* **26**, 759-774 e755 (2019).
3. Salscheider, S.L. et al. AIFM1 is a component of the mitochondrial disulfide relay that drives complex I assembly through efficient import of NDUFS5. *EMBO J* **41**, e110784 (2022).
4. Fischer, M. et al. Protein import and oxidative folding in the mitochondrial intermembrane space of intact mammalian cells. *Mol Biol Cell* **24**, 2160-2170 (2013).
5. Emsley, P., Lohkamp, B., Scott, W.G. & Cowtan, K. Features and development of Coot. *Acta Crystallogr D Biol Crystallogr* **66**, 486-501 (2010).
6. Punjani, A., Rubinstein, J.L., Fleet, D.J. & Brubaker, M.A. cryoSPARC: algorithms for rapid unsupervised cryo-EM structure determination. *Nat Methods* **14**, 290-296 (2017).
7. Liebschner, D. et al. Macromolecular structure determination using X-rays, neutrons and electrons: recent developments in Phenix. *Acta Crystallogr D Struct Biol* **75**, 861-877 (2019).
8. Ferreira, P. et al. Structural insights into the coenzyme mediated monomer-dimer transition of the pro-apoptotic apoptosis inducing factor. *Biochemistry* **53**, 4204-4215 (2014).
