## Supplementary material for "Interaction with AK2A links AIFM1 to cellular energy metabolism": Structure table

| | AIFM1 dimer | AIFM-AK2 | AIFM1-MIA40 |
|---|---|---|---|
| PDB | | | |
| EMDB | | | |
| **Data Collection and Processing** | | | |
| Microscope | Titan Krios G4i | Titan Krios G4i | Titan Krios G3i |
| Voltage (keV) | 300 | 300 | 300 |
| Magnification | 120,000 | 120,000 | 120,000 |
| Pixel size at detector (Å/pixel) | 0.46 | 0.58 | 0.654 |
| Total electron exposure ($e^{-}/Å^{2}$) | 50 | 50 | 50.82/50.57 |
| Number of frames | 468 | 468 | 48 |
| Defocus range (µm) | 0.7-1.7 | 0.7-1.7 | 0.6-2.6 |
| Automation software | EPU | EPU | EPU |
| Energy filter | Selectris | Selectris | - |
| Micrographs collected (no.) | 15,44 | 4542 | 1907/2084 |
| **For each reconstruction:** | | | |
| Final particles (no.) | 227,866 | 307,496 | 291,656 |
| Space group | P1 | P1 | P1 |
| Map sharpening B factor ($Å^{2}$) | 124.9 | 97.7 | 88.2 |
| Resolution Estimates (Å) | | | |
| FSC 0.5 (unmasked/masked) | 3.1/2.91 | 2.87/2.72 | 2.98/2.68 |
| FSC 0.143 (unmasked/masked) | 2.79/2.75 | 2.6/2.56 | 2.37/2.34 |
| FSC 0 (unmasked/masked) | 2.76/2.72 | 2.57/2.54 | 2.34/2.3 |
| **Model composition** | | | |
| Chains | 5 | 7 | 7 |
| Atoms | 7068 (Hydrogens: 0) | 7224 (Hydrogens: 0) | 7336 (Hydrogens: 0) |
| Residues | Protein: 887<br>Nucleotide: 0 | Protein: 905<br>Nucleotide: 0 | Protein: 915<br>Nucleotide: 0 |
| Water | 12 | 24 | 17 |
| Ligands | FAD: 2 | FAD: 2 | FAD: 2 |
| | NAD: 2 | NAD: 2 | NAD: 2 |
| **Model Refinement** | | | |
| **Bonds (RMSD)** | | | |
| Length (Å) (# > 4σ) | 0.005 (0) | 0.003 (0) | 0.002 (0) |
| Angles (°) (# > 4σ) | 0.591 (0) | 0.527 (0) | 0.499 (0) |
| MolProbity score | 1.50 | 1.48 | 1.61 |
| Clash score | 4.39 | 4.30 | 6.35 |
| **Ramachandran plot (%)** | | | |
| Outliers | 0.00 | 0.00 | 0.00 |
| Allowed | 4.10 | 3.92 | 3.88 |
| Favored | 95.90 | 96.08 | 96.12 |

| | | | |
|---|---|---|---|
| **Rama-Z (Ramachandran plot Z-score, RMSD)** | | | |
| whole (N = 893) | -0.40 (0.28) | -0.17 (0.28) | 0.40 (0.28) |
| helix (N = 247) | 1.35 (0.35) | 1.16 (0.35) | 1.78 (0.34) |
| sheet (N = 214) | -0.33 (0.36) | 0.63 (0.35) | 0.09 (0.34) |
| loop (N = 432) | -1.00 (0.29) | -1.30 (0.27) | -0.42 (0.30) |
| Rotamer outliers (%) | 0.28 | 0.00 | 0.00 |
| Cβ outliers (%) | NA | NA | NA |
| CaBLAM outliers (%) | 2.18 | 1.70 | 2.02 |
| ADP (B-factors) | | | |
| Iso/Aniso (#) | 7068/0 | 7224/0 | 7336/0 |
| min/max/mean | | | |
| Protein | 9.09/132.73/63.11 | 6.14/105.04/36.33 | 22.79/135.12/64.32 |
| Nucleotide | --- | --- | --- |
| Ligand | 18.37/80.80/37.07 | 8.74/40.20/21.01 | 31.78/68.66/46.21 |
| Water | 18.05/53.20/30.17 | 12.12/31.41/19.63 | 30.22/58.08/46.41 |
